# Rab5 and Alsin regulate stress-activated cytoprotective signaling on mitochondria

**DOI:** 10.1101/200428

**Authors:** FoSheng Hsu, Stephanie Spannl, Charles Ferguson, Tony Hyman, Robert G. Parton, Marino Zerial

## Abstract

Mitochondrial stress response is essential for cell survival, and damaged mitochondria are a hallmark of neurodegenerative diseases. It is thus fundamental to understand how mitochondria relay information within the cell. Here, by investigating mitochondrial-endosome contact sites we made the surprising observation that the small GTPase Rab5 translocates from early endosomes to the outer mitochondrial membrane upon oxidative stress. This is accompanied by an increase in Rab5-positive endosomes in contact with mitochondria. Interestingly, activation of Rab5 on mitochondria depend on the Rab5-GEF ALS2/Alsin, which is encoded by a gene mutated in amyotrophic lateral sclerosis (ALS). Alsin^-/-^ human induced pluripotent stem cell-derived spinal motor neurons cannot relocate Rab5 to mitochondria and display increased susceptibility to oxidative stress. These findings define a novel pathway whereby Alsin catalyzes assembly of the Rab5 endocytic machinery on mitochondria. Defects in stress-sensing by endosomes could be crucial for mitochondrial quality control during the onset of ALS.

## Introduction

Mitochondria, the energy “powerhouse” of the cell, play an essential role in a number of other cellular processes such as calcium signaling, lipid synthesis and trafficking, metabolite transport, apoptosis, and reactive oxygen species (ROS) production (Mesmin, 2016, Ott, Gogvadze et al., 2007, Rizzuto, De Stefani et al., 2012). Many of these processes necessitate communication with other cellular compartments. For example, membrane contact sites (MCS) between endoplasmic reticulum and mitochondria are important for Ca^2+^ and lipid transfer (de Brito & Scorrano, 2008), mitochondria fission (Friedman, Lackner et al., 2011), and regulation of apoptosis (Prudent, Zunino et al., 2015). Lipid droplets and peroxisomes interact with mitochondria to regulate fatty acid oxidation (Cohen, Klug et al., 2014, Pu, Ha et al., 2011). Selected membranes can be delivered to peroxisomes and lysosomes via mitochondrial-derived vesicles (Sugiura, McLelland et al., 2014). These examples demonstrate an extensive functional interplay between organelles, either directly via MCS and/or indirectly via vesicular intermediates. However, the underlying molecular mechanisms remain poorly understood and, in particular, the functional relationship between mitochondria and the endocytic system is largely unexplored.

The endocytic pathway is responsible for maintaining cellular homeostasis by internalizing, sorting, recycling and/or degrading distinct types of cargo molecules (Huotari & Helenius, 2011). Rab GTPases serve as molecular signatures for the endosomes, regulating their biogenesis and functions (Pfeffer, 2017, Zerial & McBride, 2001, Zhen & Stenmark, 2015). Ligand-receptor complexes at the plasma membrane (PM) are internalized into <u>e</u>arly <u>e</u>ndosomes (EE) marked by small GTPase Rab5, followed by either their recycling back to the PM via Rab4 and Rab11-positive recycling <u>e</u>ndosomes (RE), or conversion into Rab7-positive late <u>e</u>ndosomes (LE) *en route* to lysosomes for degradation (Rink, Ghigo et al., 2005, Sonnichsen, De Renzis et al., 2000, Ullrich, Reinsch et al., 1996). On these endosomal compartments, Rab proteins recruit a plethora of effectors for membrane tethering and fusion, cargo sorting and signaling (Sorkin & von Zastrow, 2009, Stenmark, 2009, Zerial & McBride, 2001). For example, EEA1 is a dimeric coiled-coiled RAb5 effector protein that tethers two vesicles to allow efficient fusion between Rab5-harbouring membranes (Murray, Jahnel et al., 2016). Other Rab5 effectors such as APPL1 are involved in regulating metabolic and inflammatory responses (Schenck, Goto-Silva et al., 2008, Wen, Yang et al., 2010). Rab activation, and thus stabilization after recruitment, on the membrane requires guanine nucleotide exchange factors (GEFs) (Blumer, Rey et al., 2013). In the case of Rab5, GEFs constitute a family of VPS9 domain-containing proteins, including Rabex-5 (Horiuchi, Lippe et al., 1997), RME-6 (Sato, Sato et al., 2005), amyotrophic lateral sclerosis protein 2 (ALS2/Alsin) (Otomo, Hadano et al., 2003), and mammalian Ras and Rab interactor 1, 2, 3 (Rin1-3) (Hu, Bliss et al., 2005). The rationale behind this complexity is that Rab5 must be specifically regulated by the different GEFs in space and time. The rationale behind this complexity is that Rab5 must be specifically regulated by the different GEFs in space and time. In this respect, the function of many Rab5 GEFs remains unclear.

Physical interactions between the endosomal machinery and mitochondria serve important functions in cell homeostasis, repair and apoptosis. For example, transfer of iron (Das, Nag et al., 2016, Sheftel, Zhang et al., 2007) and cholesterol (Charman, Kennedy et al., 2010) from endosomes to mitochondria is enabled by physical interactions between the two organelles. Another classical example of mitochondria-endo-lysosome interactions is autophagy. Autophagy is a clearance mechanism whereby cells identify defective organelles following damage or stress and eliminate them via the formation of an autophagosome and fusion with lysosomes (Mizushima & Levine, 2010). The mechanism of degrading mitochondria has been termed macroautophagy or mitophagy. Intriguingly, expression of pro-apoptotic factors such as canonical BH3-only proteins drive Rab5- and Rab7-positive endolysosomes into inner mitochondrial compartments, via a pathway that appears to differ from autophagy/mitophagy (Hamacher-Brady, Choe et al., 2014). Interestingly, our previously conducted genome-wide RNAi screen of endocytosis (Collinet, Stoter et al., 2010) revealed that ∼8% of the hit genes had mitochondrial-related functions, pointing at hitherto unexplored molecular connections between the endosomal system and mitochondria. This led us to hypothesize that other mitochondrial functions may be regulated by endocytic components.

Here, by exploring interactions between early endosomes and mitochondria, we made an unexpected observation: we found that upon laser- or chemically-induced oxidative stress in mammalian cells, mitochondria outer membrane permeabilization (MOMP) releases mitochondrial factors such as cytochrome c, and concomitantly, triggers the assembly of the Rab5 machinery on the OMM, in a process independent of mitophagy. Remarkably, we found that the Rab5 GEF responsible for Rab5 activation is Alsin/ALS2, which is also recruited to OMM. Our findings suggest that the Rab endocytic machineries intimately interact with mitochondria during oxidative stress as a cytoprotection mechanism with important implications for amyotrophic lateral sclerosis (ALS) and other neurodegenerative diseases.

## Results

### Inter-organelle contacts between endosomes and mitochondria

We first explored the potential physical link between the endosomal system and mitochondria at steady state. In order to capture early endocytic events, HeLa cells stably expressing TagRFP-MTS (mitochondria targeting sequence) (Takeuchi, Kim et al., 2013) were incubated with endocytic cargoes such as Alexa-conjugated transferrin (Tfn) or epidermal growth factor (EGF) for 10 min at 37°C. Cells were fixed and imaged via confocal microscopy. All acquired images were subjected to chromatic shift correction, deconvolution, and localization analysis (MotionTracking) based on subtraction of random colocalization (Kalaidzidis, 2015). Both Tfn and EGF were consistently observed in a subset of endosomes that partially overlapped or were in close proximity to mitochondria (Figure 1A and B). Given that endosomes are motile and omnipresent in the cytoplasm, and the resolution limit of conventional light microscopy, it is difficult to determine whether the observed proximity of endosomes to mitochondria reflects real MCS or simply due to random chance (Kalaidzidis, 2015). One way to approach this problem is to explore how dynamic such inter-organelle contacts are during early endocytic events. To assess the interactions, cells were incubated with Tfn-488 for 1 min to label early endosomes and immediately imaged live using a spinning disk confocal microscope. In the 10-min time-lapsed videos (e.g. Video 1), some organelle interactions were visible between Tfn-containing endosomes and mitochondria labelled by TagRFP-MTS. Endosomal vesicles remained in the proximity or in contact with mitochondria for 3-5 min and, interestingly, we could observe interactions that were followed by fission-like events (Video 2). The live-cell imaging results suggest that the proximity of endosomes and mitochondria observed at steady state (Figure 1) may reflect *bona fide* albeit transient interactions, as suggested previously (Das, Nag et al., 2016, Sheftel, Zhang et al., 2007).

**Figure 1.**
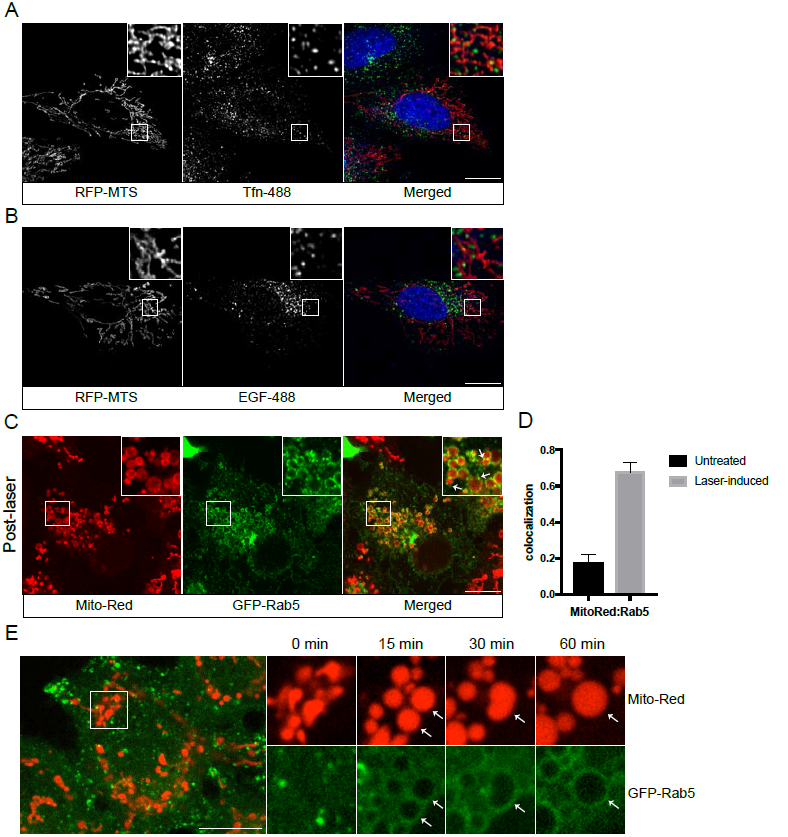
Recruitment of Rab5 to the outer mitochondrial membrane (OMM) upon mitochondrial stress. (**A**) and (**B**) HeLa cells were transfected with TagRFP-MTS (mitochondria targeting sequence) and labeled with either Alexa-488 transferrin (Tfn) or epidermal growth factor (EGF), respectively. (**C**) Live BAC GFP-Rab5 HeLa cells were labeled with 100 nM MitoTracker-Red CMXRos for 30 min at 37°C. Cells were photoirradiated with low dosage of 561 nm laser (∼5 J/cm^2^) for 60 sec. Snapshot of the cell was taken at 15 min post-laser treatment. Arrowheads indicate GFP-Rab5-positive endosomes that are in close proximity to mitochondria. (**D**) Quantification of colocalization before (Untreated) and post-laser treatment (Laser-induced), *n* = 3. The y-axis is defined as the ratio of the amount of MitoTracker-Red colocalized with GFP-Rab5 to the total amount of MitoTracker-Red. Error bars represent SEMs. (**E**) Time-lapse images of a cell following laser treatment. Inset images are shown at different time points (min). Arrowheads indicate recruitment of GFP-Rab5 to the mitochondria within 15 min, which persist over a period of 60 min. Scale bars, 10 μm.

### Acute mitochondrial stress recruits Rab5 and Rab5-positive endosomes to the outer mitochondrial membrane

Given the key role of mitochondria in sensing and responding to oxidative stress, we asked whether acute perturbation of mitochondria affects endosomes-mitochondria interactions. We used HeLa cells stably expressing GFP-Rab5 under its endogenous promoter with a bacterial artificial chromosome (BAC) transgene (BAC GFP-Rab5) (Villasenor, Nonaka et al., 2015). Live-cell imaging of BAC GFP-Rab5 expressing RFP-MTS frequently confirmed the presence of Rab5-positive early endosomes (>200 nm) in close contact with mitochondria (Video 3). In addition to using an ectopically expressed mitochondrial marker, we also tested other mitochondrial-selective dyes in our live-cell imaging experiments. Unexpectedly, we found that upon prolonged illumination, there was not only a change in mitochondrial morphology but also an alteration in GFP-Rab5 dynamics, with varying levels of increased signal around mitochondria depending on the nature of the dyes. Certain rosamines and rhodamine-derived dyes used to assay mitochondrial functions possess photosensitizing properties (Hsieh, Chu et al., 2015). Therefore, we hypothesized that the change in Rab5 dynamics may be a direct consequence of the change in mitochondrial function.

To this end, we applied MitoTracker-Red CMXRos to specifically perturb mitochondrial function (Minamikawa, Sriratana et al., 1999). Consistent with previous reports, low dosage with 561 nm laser (∼5 J/cm^2^) caused a decrease in mitochondrial and an increase in cytoplasmic signal, indicative of MOMP, accompanied by globular swelling of mitochondria within min (Figure 1C, Video 4). Surprisingly, laser treatment on MitoTracker-Red labeled cells resulted in translocation of GFP-Rab5 to OMM, marked by an increased in co-localization compared to untreated (Figure 1D). Line scan analysis showed that the recruitment of Rab5 to mitochondria occurred within min post laser treatment (Figure 1–figure supplement 1A). The Rab5 ring-like signal formed within min and sustained for >60 min where fusion between two mitochondria can be seen (Figure 1E, arrowheads). We also frequently observed the presence of endosomes in proximity to the altered mitochondria (Figure 1C, arrowheads). As controls, cells labeled with MitoTracker Green FM, which does not share the photosensitizing properties reported for MitoTracker-Red CMXRos, or transfected with RFP-MTS retained their tubular structures under the same laser treatment (Figure 1–figure supplement 1B,C), consistent with previous findings (Minamikawa et al., 1999). Additionally, we tested the specificity and localization of the GFP signals by Rab5 and outer mitochondrial membrane protein TOM20 antibodies (Figure 1–figure supplement 2). These results suggest that Rab5 translocates to mitochondria upon MOMP.

### Rab5 localizes to regions of mitochondria that are damaged

We next asked whether GFP-Rab5 relocalization to mitochondria is a general response to overall cell stress or can be elicited locally on individual mitochondria. To test this, we applied stress in a small area within a cell labeled with MitoTracker-Red (Figure 2A, Pre, inset) and monitored the GFP-Rab5 signal after 10 and 20 min (Figure 2B, Post-laser). At 10-minute time after laser irradiation, the MitoTracker-Red signal began to fade. Despite the localized perturbation, we observed an overall change in mitochondrial morphology, from tubular to rounded. This is consistent with the fact that mitochondria form a dynamically interconnected network (Lackner, 2014, Wang, Du et al., 2015) that appears to react to local damage as an ensemble. However, at 20-minute time point, we found strong Rab5 recruitment only in the laser-induced area and not in the rest of the cell (Figure 2B). In the laser-induced area, Rab5 rings formed on rounded mitochondria that exhibited a loss in luminal MitoTracker dye, presumably as a result of MOMP (Figure 2B, arrowheads). We also observed the appearance of distinct Rab5-positive endosomes contacting these mitochondria (Figure 2B, double arrowheads). These results suggest that Rab5 is recruited in response to signal(s) originated from individually damaged mitochondria.

**Figure 2.**
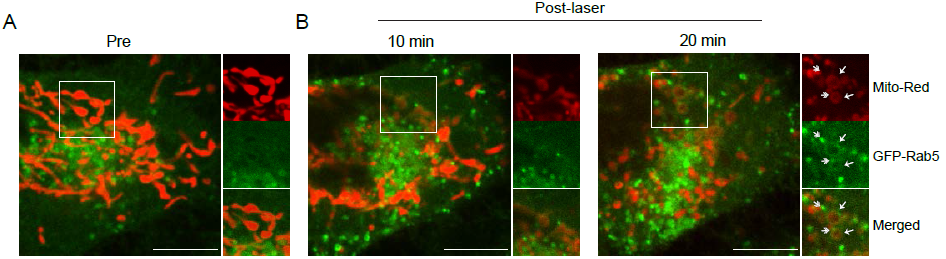
Rab5 is signaled onto localized damaged mitochondria. (**A**) BAC GFP-Rab5 cells were labeled with 100 nM MitoTracker-Red CMXRos for 30 min at 37°C, and imaged live. A cell before laser treatment is shown (Pre). (**B**) Photoirradiation was applied as before to a localized region within the cell (white box). Snapshot of the cell was taken at 10 min and 20 min post-laser. Inset images indicate the effect of laser treatment on mitochondrial morphology and network. Arrowheads indicate the recruitment of GFP-Rab5 (filled) and GFP-Rab5-positive endosomes near the OMM (double-line) at 20 min time point. Scale bars, 10 μm.

### Membrane contacts between Rab5-positive mitochondria and endosomes

In our stress-induced conditions, we frequently observed Rab5-positive endosomes in close proximity to the swollen mitochondria (Figure 1C, Figure 2B, double arrowheads, Figure 1–figure supplement 1A). By live-cell imaging, these endosomes also appeared to dock stably onto the mitochondria (Video 5, boxed regions). Due to the diffraction limit of standard light microscopy, we could not resolve objects that are closer than 200 nm and endosomes that are <200 nm in diameter. For these reasons, we complemented our study by correlative light and electron microscopy (CLEM) to obtain ultrastructural details. We designed our experimental setup to 1) image an entire cell live, 2) visualize the translocation of GFP-Rab5 and Rab5-positive endosomes onto mitochondria upon laser-induced stress, and 3) re-image the same cell by serial section transmission electron microscopy (TEM) (Figure 3A). GFP-Rab5 cells labeled with MitoTracker-Red CMXRos were plated on gridded culture dishes, laser-treated and imaged live on the spinning disk confocal microscope. Upon mitochondrial rounding, cells were immediately fixed and re-imaged again to obtain post-fixation image. These showed distinct GFP-positive puncta in close proximity to mitochondria (Figure 3A,B, red arrowhead). Samples were then processed for serial section transmission electron microscopy (TEM) and the same region was re-located in the thin sections by both nuclear membrane (Figure 3B, dotted line) and mitochondria (Figure 3B, red star: mitochondria), acting as fiducial markers. The TEM images from three different serial sections of the inset area in Figure 3B revealed that the GFP-Rab5-positive structure corresponds to a tubular-cisternal structure, approximately 400nm in diameter, with the typical morphology of an early endosome (Figure 3C, green). Serial section analysis showed that the endosomal membrane was in very close contact (<5 nm) with the adjacent mitochondrion (Figure 3C, red). The rounded mitochondrion showed few cristae, a diameter of approximately 1.5 μm, and lacked any additional enveloping membranous structures that might be indicative of mitophagy (Youle & Narendra, 2011). Our data suggest that upon mitochondrial stress, two events involving early endosomes and mitochondria occur: 1) Rab5 translocates to mitochondria and 2) endosomes and mitochondria engage in membrane contacts.

**Figure 3.**
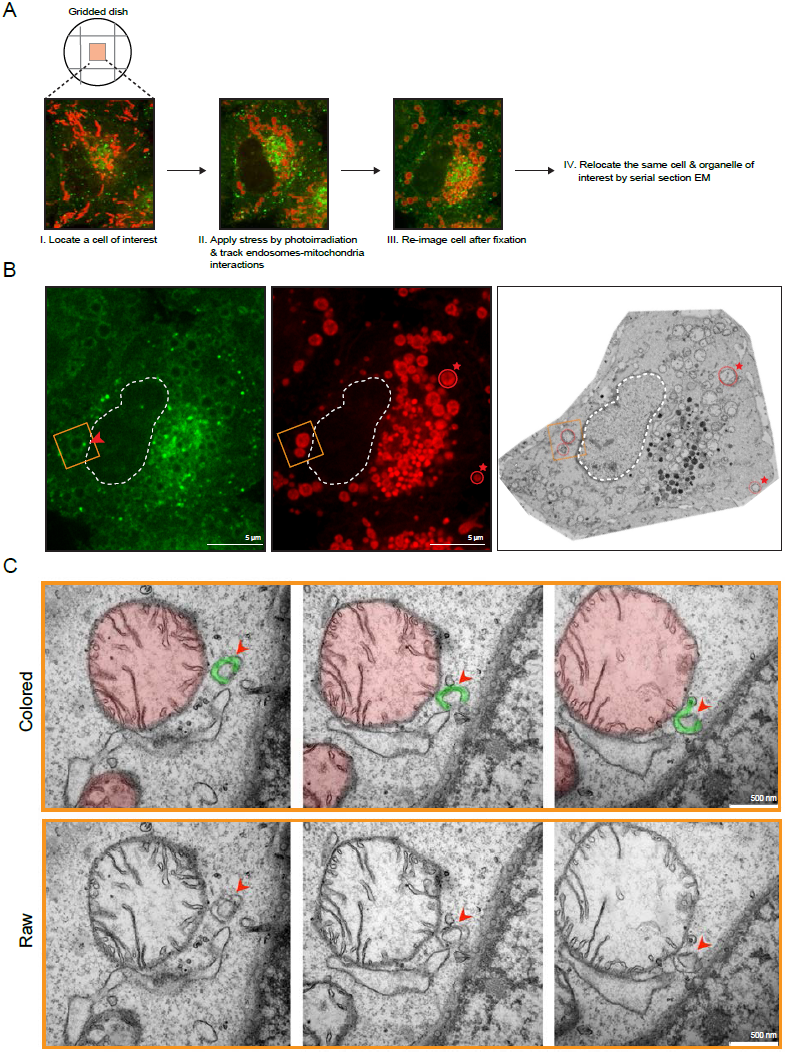
Membrane contacts between Rab5-positive mitochondria and endosomes. Ultrastructural analysis of cells upon laser-induced stress. (**A**) Flow chart of the experimental set up. BAC GFP-Rab5 cells were seeded onto gridded dish, labeled with MitoTracker-Red CMXRos. The motility of GFP-Rab5-positive endosomes were briefly assessed as a readout for healthy and active cells. A cell of interest was then photoirradiated with lower dosage of 561 nm laser. (**B**) Fluorescence images of GFP-Rab5 and MitoTracker-Red after fixation with glutaraldehyde. Box region shows a GFP-Rab5 endosome (arrowhead) next to a swollen mitochondrion. The transmission electron micrograph (TEM) of the same cell is shown. The nuclear membrane (dotted line) and mitochondria (red star) were used as fiducial markers to re-locate the same cell. (**C**) Zoom-in images of the box region in (**B**). Colored images indicate GFP-Rab5-positive structure in green and mitochondria in red. Raw images are shown below. Mitochondrial cristae are visible.

### The recruitment of Rab5 to mitochondria is not due to mitophagy

Following mitochondrial stress, both the timing (in min) of Rab5 recruitment on the rounded mitochondria and the absence of wrapped double-membrane structures argue against mitophagy (Narendra, Tanaka et al., 2008, Novak, Kirkin et al., 2010). We searched for additional evidence to rule out this mechanism. Mitophagy requires the E3 ubiquitin ligase Parkin (Narendra et al., 2008). Parkin is normally located in the cytosol but is recruited to damaged mitochondria, followed by formation of LC3-positive autophagosomes and fusion with Lamp1-positive lysosomes in a process that occurs in hours to days (Dolman, Chambers et al., 2013). To test whether the swollen mitochondria observed in our system undergo this process, we examined the localization of the three markers in BAC GFP-Rab5 cells. We first labeled live cells with MitoTracker-Red CMXRos in order to image mitochondria and endosomes at steady state (Figure 4A,C,E, Untreated) prior to triggering laser-induced stress as before. We then incubated the cells for 60 min as a mean to maximize the time window that these markers might be recruited. Upon fixation, cells were immunostained with specific antibodies to detect endogenous LC3, Lamp1, and Parkin. Since cells were seeded on a gridded dish as positional marker, we were able to re-locate the same cell, which allowed us to assess any changes to the localization of the different markers as a result of stress when compared to neighboring untreated cells (Figure 4–figure supplement 1). In all laser-induced cells, GFP-Rab5 was specifically enriched around mitochondria when compared to untreated (Figure 4B, D, and F). Sparse LC3 puncta were observed near the perinuclear region in either neighboring untreated or laser-treated cells, where the majority of mitochondria were devoid of any LC3 signals (Figure 4B,G, Figure 4–figure supplement 1A). Similarly, Lamp1 puncta, which mainly resided in the perinuclear region, did not show any change in location in neither untreated nor laser-treated cells (Figure 4D,G, Figure 4–figure supplement 1B). Parkin also remained cytoplasmic and did not show any enrichment around mitochondria upon laser treatment in the time frame of our experiment (Figure 4F,G, Figure 4–figure supplement 1C).

**Figure 4.**
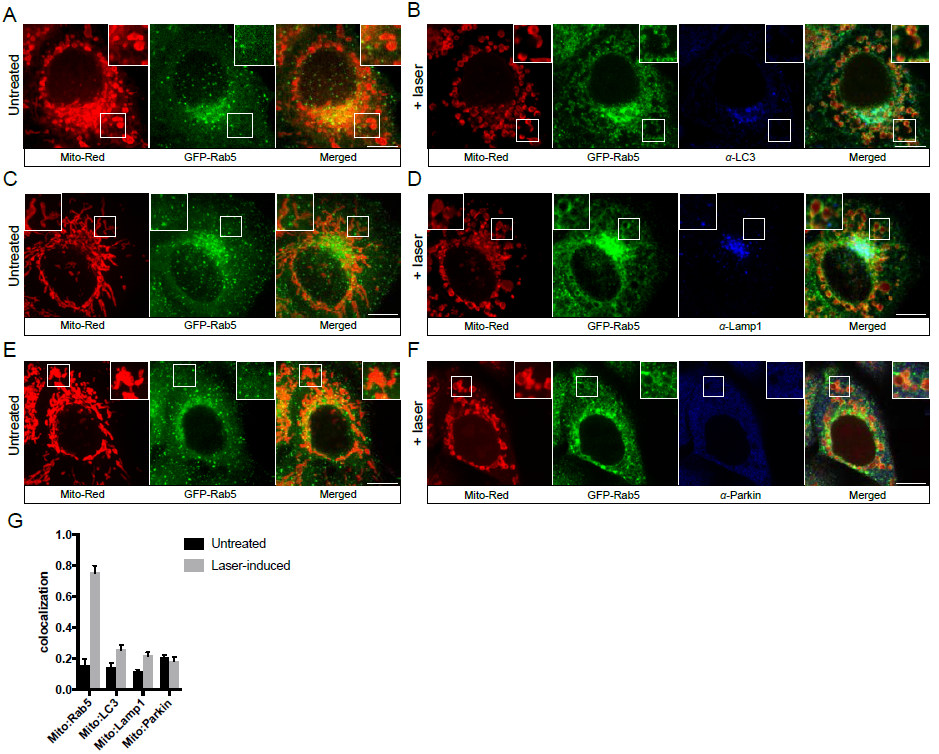
Localization of endogenous LC3, Lamp1, and Parkin upon mitochondrial stress. BAC GFP-Rab5 cells were labeled with 100 nm of MitoTracker-Red CMXRos for 30 min at 37°C. Snapshot images of cells were taken as control (Untreated) (**A**), (**C**), and (**E**). The same cells were photoirradiated as before, fixed after 60-min elapsed, and labeled with specific antibodies against LC3 (**B**), Lamp1 (**D**), and Parkin (**F**), respectively. Inset images indicate the approximate area before and after laser treatment. (**G**) Quantification of colocalization of MitoTracker-Red to Rab5, LC3, Lamp1, and Parkin, respectively, in untreated versus laser-induced conditions (*n* = 3). Untreated cells correspond to cells outside of laser treated area. Scale bars, 10 μm.

In addition to examining endogenous proteins by immunostaining, we also tested all three markers by live-cell imaging in HeLa BAC cell lines expressing GFP-tagged LC3, Lamp1, and Parkin (Figure 4–figure supplement 2). Cells were laser-induced as before and monitored live for 60 min. As observed with endogenous LC3, a fraction of GFP-LC3 was enriched in the perinuclear region, where it overlapped with small fragmented mitochondria, but not the rest of the mitochondria (Figure 4 – figure supplement 2A). We also observed weak GFP-Parkin recruitment to small fragmented mitochondria whereas most mitochondria were devoid of signal in laser-induced conditions (Figure 4 – figure supplement 2C, arrowheads). The enrichment to these small fragmented mitochondria may be a result of over-expression which activates some level of mitophagy (Rana, Rera et al., 2013). Nevertheless, unlike Rab5, Parkin was not recruited to the majority of mitochondria. Altogether, the kinetics of Rab5 to stressed mitochondria is indicative of a very fast response (<10 min), the absence of a double membranous structure (Figure 3C) and the lack of significant Parkin recruitment post 3 h treatment (data not shown) argue that the translocation of Rab5 to mitochondria occurs much earlier than the onset of autophagy and mitophagy.

### Rab5 translocation to mitochondria is linked to release of cytochrome c upon hydrogen peroxide treatment

What could be the signal that drives Rab5 recruitment? Several possible scenarios such as morphological changes to mitochondria and/or release of mitochondrial-derived factor(s) may be accounted for. Morphological changes such as matrix condensation or swelling of mitochondria are often associated with MOMP, cytochrome c release, and subsequent activation of caspases (Gottlieb, Armour et al., 2003). However, this is not a prerequisite. For example, the protonophore carbonyl cyanide m-chlorophenyl hydrazone (CCCP) causes mitochondrial swelling and rounding without immediate cytochrome c release nor cell death (Gao, Pu et al., 2001, Lim, Minamikawa et al., 2001). On the other hand, hydrogen peroxide (H_2_O_2_) was reported to induce mitochondrial rounding associated with cytochrome c release and caspase activation (Takeyama, Miki et al., 2002). To address this question, we tested CCCP and H_2_O_2_ on the effect of Rab5 localization. In DMSO control cells, mitochondria were mostly tubular (Figure 5A,B, top panels). The exposure of cells to either CCCP or H_2_O_2_ for 2 h resulted in mitochondrial rounding, similar to the effect with laser-treatment (Figure 5A,B, bottom panels). Interestingly, Rab5 enrichment on mitochondria was only observed in H_2_O_2_-treated cells and not in CCCP-treated cells (Figure 5A,B, arrowheads). The enrichment of Rab5 to mitochondria in H_2_O_2_ condition was ∼4-fold higher compared to control cells, as revealed by co-localization analysis (Figure 5C,D).

**Figure 5.**
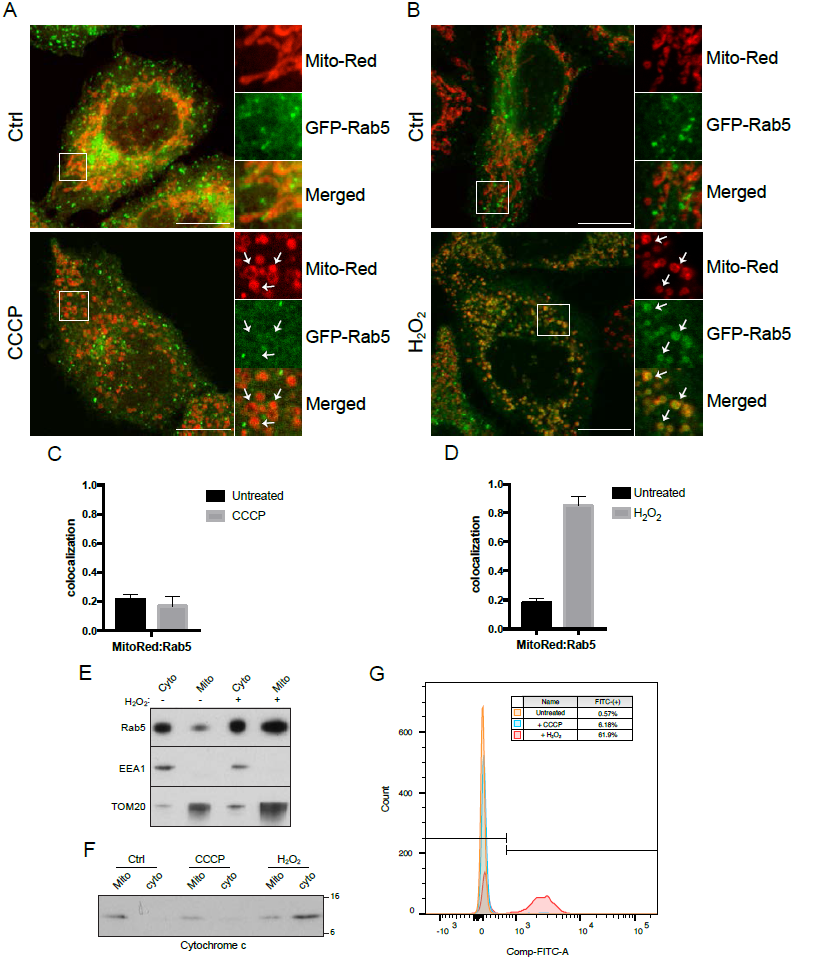
Effects of carbonyl cyanide m-chlorophenyl hydrazone (CCCP) and hydrogen peroxide (H_2_O_2_) on Rab5 recruitment to mitochondria. (**A**) Cells were treated with either DMSO (Ctrl) or 10 μM CCCP for 2 h. (**B**) Cell were treated with either DMSO (Ctrl) or 250 μM H2O_2_ for 2 h. Cells were fixed and imaged by confocal microscopy. Inset regions show the effect of the treatment on mitochondrial morphology, and GFP-Rab5 localization. Arrowheads point to rounded mitochondria in both CCCP- and H_2_O_2_-treated conditions. (**C**) and (**D**) Quantification of colocalization in (**A**) and (**B**), respectively. *n* = 10. resulted in robust recruitment of GFP-Rab5. (**E**) Subcellular fractionation of cytosolic (Cyto) and mitochondrial (Mito) fractions from HeLa cells treated with either PBS (control) or 250 μM H2O_2_ for 2 h at 37°C. Protein samples were loaded onto SDS-PAGE and immunoblotted with Rab5, EEA1, and TOM20 antibodies. (**F**) Subcellular fractionation of HeLa cells treated with DMSO, 10 μM CCCP, or 250 μM H2O_2_ for 2 h. Protein samples from purified mitochondria (Mito) and cytosolic (Cyto) fractions were loaded onto SDS-PAGE and imunoblotted with cytochrome c antibody by Western blot. (**G**) Cells were treated the same way as in (**F**). Cells were then resuspended in live-cell imaging solution containing 500 nM caspase-3/7 green flow cytometric reagent and incubated for 30 min at 37°C before subjecting to flow cytometric analysis. FITC signal (x-axis) is plotted against total cell count (y-axis). Gating was set based on background signal in DMSO control. Error bars represent SEMs.

To corroborate these morphological observations with an independent method, we also tested the effect of H_2_O_2_ on Rab5 association with endosomes and mitochondria by subcellular fractionation. We isolated cytosolic and mitochondrial fractions via differential centrifugation and assessed the purity by Western blot analysis using TOM20 as mitochondrial marker and EEA1 as endosomal marker. Consistent with the observation that Rab5 is translocated to stressed mitochondria, we found that cells challenged with H_2_O_2_ showed a strong increase in the amount of Rab5 present in the mitochondrial fraction (Figure 5E).

We then asked whether the observed difference between CCCP and H_2_O_2_ could be related to the release of mitochondrial factors such as cytochrome c. To this end, we performed subcellular fractionation on cells treated with either 10 μM CCCP or 250 μM H_2_O_2_ for 2 h. The release of cytochrome c into the cytosol was upregulated in H_2_O_2_-treated cells, but not in CCCP-treated cells (Figure 5F), consistent with previous reports (Takeyama et al., 2002). Since cytochrome c is a known factor for activating caspase-dependent programmed cell death, we assessed the activity of caspase 3/7 via a 4-amino acid peptide (DEVD) conjugated to a DNA-binding dye. Cleavage of the DEVD peptide by caspase 3/7 releases the DNA-binding fragment, yielding a fluorescent signal. Using flow cytometry, we detected ∼62% of cells showing strong fluorescent signal in H_2_O_2_-treated cells, and merely ∼0.6% and ∼6.2% in control and CCCP-treated, respectively (Figure 5G).

Altogether, these results show that, in addition to the morphological alterations, the release of cytochrome c and consequent activation of the caspase-dependent apoptotic pathway, H_2_O_2_ treatment also induces the translocation of Rab5 to mitochondria.

### Rab5 translocation to mitochondria blocks caspase release and inhibits trafficking of transferrin upon oxidative stress

In the course of H_2_O_2_ treatment by live-cell imaging, we found that mitochondria appeared to respond with different kinetics within an individual cell (Video 6). At the 60-min time point, distinct regions of the mitochondrial network were more prone to rounding and membrane permeabilization than others, as revealed by the differential loss of MitoTracker-Red signal when compared to control at 0 min (Figure 6A, inset image). Interestingly, these regions correlated exclusively with Rab5 ring-like recruitment (Figure 6A, inset image, arrowheads). This suggests that Rab5 may be involved in either facilitating or preventing the apoptotic process. To address this, we over-expressed either GFP or GFP-Rab5 in HeLa cells and measured the amount of cytosolic cytochrome c at different time points after H_2_O_2_ addition. We found a significant delay in cytochrome c release from mitochondria in GFP-Rab5 transfected cells compared to control cells (Figure 6B,C). These results suggest that Rab5 plays a protective role in mitochondrial-induced apoptosis by down-regulating the release of pro-apoptotic factors into cytosol.

**Figure 6.**
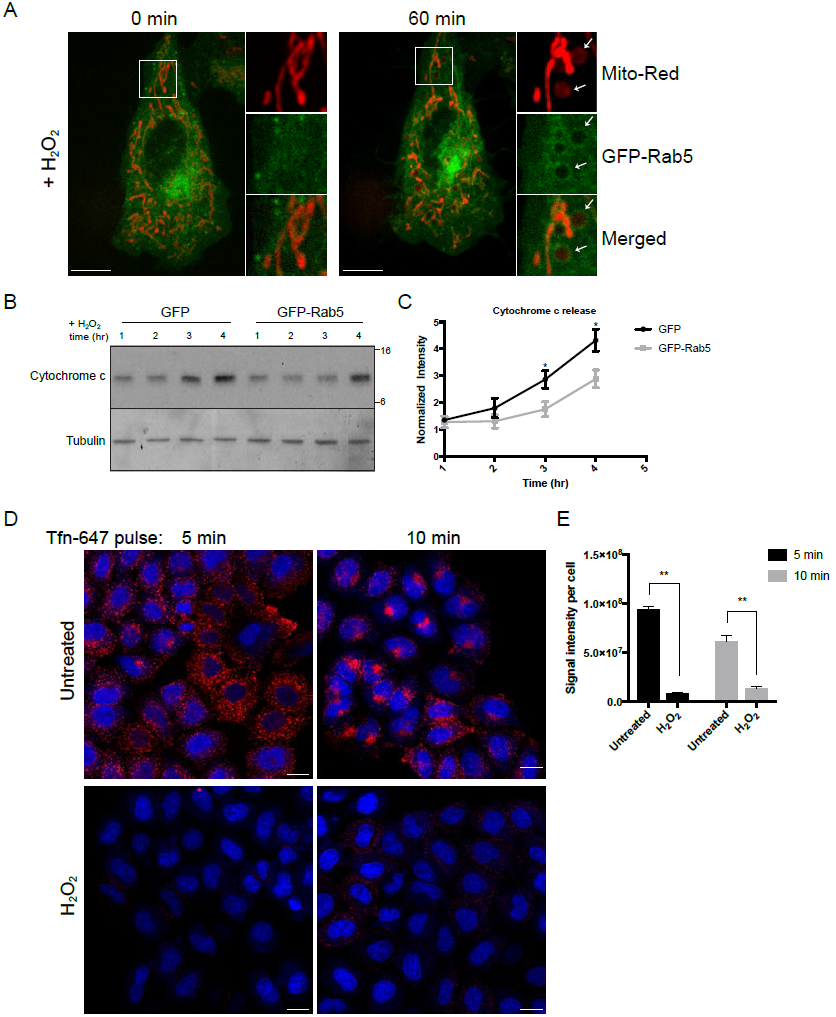
Rab5 regulates mitochondrial-induced apoptosis by blocking cytochrome c release. (**A**) Representative images of a live BAC GFP-Rab5 cell before (0 min) and after treatment with 250 μM H_2_O_2_ for 60 min. MitoTracker-Red CMXRos was used to monitor MOMP events as shown in the inset images (right panel, arrowheads). Scale bars, 10 μm. (**B**) HeLa cells over-expressing either GFP or GFP-Rab5 were treated with 250 μM H_2_O_2_ at different time points (1hr to 4 hr). Protein samples from purified cytosolic fraction were obtained and immunoblotted with cytochrome c antibody by Western blot. Tubulin was included as loading control. (**C**) Densitometric quantification from the Western blots performed in (**B**). Data were collected from three independent experiments (*n* = 3). Y-axis corresponds to normalized ratio intensity of experimental to loading control (* p <0.05). (**D**) Cells were seeded in 384-well plate and incubated with complete medium (Untreated) or in the presence of 250 μM H_2_O_2_ for 2 hours at 37°C. Cells were then pulsed with Alexa-647 Tfn (10 ug/ml) for 5 min and 10 min, followed by PBS wash, fixed with PFA, and stained with DAPI (nuclear) and CellMask Blue (cytoplasmic). Images were acquired by the Yogokawa confocal microscope. Scale bars, 10 μm. (**E**) Quantification of cytoplasmic fluorescence intensity per cell, *n* = 30 (** p <0.005).

Given the key role of Rab5 in the biogenesis of the endosomal system (Zeigerer, Gilleron et al., 2012), the dramatic translocation of Rab5 to mitochondria upon oxidative stress by H_2_O_2_ led us to ask whether endocytic trafficking is affected. To address this, we stimulated HeLa cells with Alexa-647 Tfn continuously for 5 and 10 min. In the absence of H_2_O_2_, transferrin was present in endosomes throughout the cells and at 10 min accumulated in perinuclear recycling endosomes (Maxfield & McGraw, 2004) (Figure 6D, top). In contrast, cells treated with 250 μM H_2_O_2_ showed a severe block in Tfn accumulation at both 5 min and 10 min (Figure 6D,E). This suggests that endosomal trafficking is inhibited during mitochondrial stress, consistent with the reduction of Rab5 on endosomes and its re-location to mitochondria.

### Rab5 enrichment on the OMM is accompanied by specific effector recruitment

Since Rab5 translocates from early endosomes to mitochondria, with consequent reduction in endocytic uptake, we next asked whether endosomal Rab5 effectors are also recruited onto mitochondria. We were able to systematically assess the localization of various endosomal effectors such as Rabenosyn-5/ZFYVE20, EEA1 and APPL1/2 in BAC GFP-Rab5 HeLa cells labeled with MitoTracker-Red CMXRos via immunostaining by pair-wise combinations. We aimed at detecting the endogenous rather than the tagged proteins as the latter often cause perturbations and do nor recapitulate the native protein function (Kalaidzidis JCB 2016). The specific antibodies were first tested in untreated control cells, which all showed significant levels of co-localization with GFP-Rab5 (Figure 7–figure supplement 1). Upon laser-induced stress, the GFP-Rab5 recruitment of Rab5 rings around mitochondria provided an immediate visual cue and served as positive control. All cells were fixed after 30 min incubation post-laser treatment. Out of the effectors tested, we detected a strong enrichment of Rabenosyn-5/ZFYVE20 to mitochondria but not EEA1 in the same cell (Figure 7A,C). Neither APPL1 nor APPL2 showed enrichment around mitochondria, despite strong Rab5 ring formation (Figure 7B,C). Unlike Rabenosyn-5/ZFYVE20, EEA1 and APPL1/2 remained well distributed in endosomal-like vesicles as in both treated and untreated cells (Figure 7B, Figure 7–figure supplement 1C,D). When we assessed the co-localization of Rab5 and EEA1, there was a loss in co-localization in laser-treated compared to untreated cells (Figure 7D). The unique localization pattern led us to ask whether phosphatidylinositol 3-phosphate (PI(3)P) was on OMM in our stress conditions, since both ZFYVE20/Rabenosyn-5 and EEA1 are recruited to endosomes via both Rab5 and PI(3)P-binding FYVE motifs (Nielsen, Christoforidis et al., 2000). To test this, we over-expressed the PI(3)P probe GFP-2xFYVE^Hrs^ (Gillooly, Morrow et al., 2000) in HeLa cells and monitored the signals in live cells upon laser-induced stress. At the initial time point, mitochondria started to retract and swell up, and the GFP signals were present as vesicle-like puncta (Figure 7–figure supplement 2, 0 min) consistent with previously reported localization (Gillooly, Morrow et al., 2000). At 60-min elapsed time point, all mitochondria appeared swollen but were completely devoid of GFP signals, which remained on vesicle-like puncta (Figure 7–figure supplement 2, 60 min).

**Figure 7.**
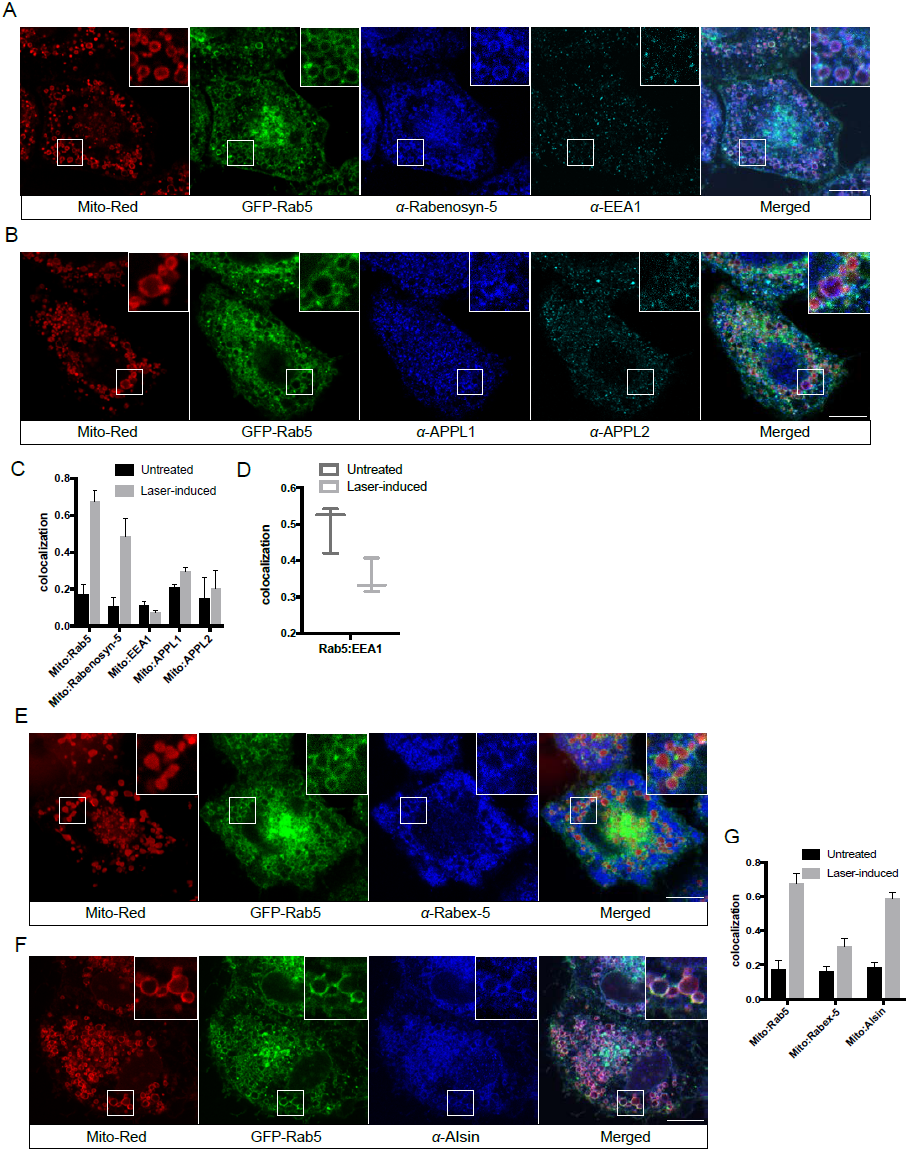
Localization of Rab5 effectors upon laser-induced mitochondrial damage. BAC GFP-Rab5 cells seeded on a gridded dish were labeled with MitoTracker-Red CMSRox and photoirradiated as before. Cells were fixed after 30 min post-laser treatment and immunostained with specific ZFYVE20 and EEA1 (**A**), and APPL1 and APPL2 antibodies (**B**). (**C**) Quantification of colocalization from untreated and laser-treated cells in (**A**) and (**B**), *n* = 3. (**D**) Quantification of colocalization between Rab5 and EEA1 as shown in (**A**), *n* = 3. (**E**-**F**) BAC GFP-Rab5 cells were treated and processed in the same manner as in (**A**) and immunostained with specific Rabex-5 and Alsin antibodies, respectively. (**G**) Quantification of colocalization from untreated and laser-treated cells in (**E**) and (**F**), *n* = 3. Scale bars, 10 μm.

Our findings reveal a selective mechanism of Rab5 translocation and activation on mitochondria that result in a PI(3)P independent recruitment of specific effectors.

### The Rab5 GEF Alsin localizes to mitochondria upon stress induction

Translocation of Rab5 and recruitment of effectors imply that Rab5 must be activated on the mitochondrial membrane. Activation of Rab GTPases on organelle membranes depends on a family of GEFs (Blumer et al., 2013, Pfeffer, 2013, Zerial & McBride, 2001, Zhen & Stenmark, 2015). We first examined the localization of Rabex-5, a known GEF of Rab5 on the endosomal membrane, by immunostaining in BAC GFP-Rab5 cells. Again, we had to localize the endogenous protein because tagged Rabex-5 constructs proved to induce artifacts on the endosomal system (Kalaidzidis and Zerial, unpublished). The formation of GFP-Rab5 rings served as positive control upon laser-treatment. Despite a slight enrichment of endogenous Rabex-5 upon laser-induced stress, the signal appeared mostly as cytosolic and cytoplasmic puncta (Figure 7E), consistent with its endosomal localization (Figure 7–figure supplement 3A).

The weak recruitment of Rabex-5 led us to hypothesize that another GEF might be principally involved. We turned our attention to Alsin as a potential candidate GEF for Rab5 on mitochondria based on several lines of evidence. Alsin is the gene product of ALS2, which is mutated in multiple neurodegenerative disorders such as juvenile amyotrophic lateral sclerosis (ALS), juvenile primary lateral sclerosis (JPLS), and infantile-onset ascending hereditary spastic paralysis (IAHSP). The gene has two splice isoforms, which encodes a long form (LF) of 1657 amino acids and a short form (SF) of 396 amino acids. Alsin-LF comprises of several GEF domains: a RCC1-like domain that acts as GEF for Ran GTPase, DH-PH domain for Rho GTPase, and a C-terminal VPS9 domain for Rab5 (Topp, Carney et al., 2005) (Figure 7–figure supplement 3B). Functional studies in ALS mouse model have associated Alsin with neuronal survival (Kanekura, Hashimoto et al., 2004, Panzeri, De Palma et al., 2006) and endolysosomal trafficking (Hadano, Mitsui et al., 2016, Hadano, Otomo et al., 2010). Moreover, corticospinal motor neuron (CSMN) in Alsin KO mice display selective defects in mitochondrial morphology (Gautam, Jara et al., 2016). We therefore tested Alsin localization under our stress-induced conditions. At steady state, Alsin was found to localize to vesicular structures, showing only partial overlap with Rab5 (Figure 7–figure supplement 3C). The staining pattern observed is consistent with the reported localization of Alsin (Kanekura et al., 2004, Topp, Gray et al., 2004). In contrast after laser treatment we found strong and uniform Alsin staining around mitochondria (Figure 7F), which showed significant co-localization with mitochondria (Figure 7G). Our data point Alsin as a candidate GEF for activating Rab5 on OMM upon stress induction.

### Alsin regulates mitochondrial apoptotic signaling and is required for efficient Rab5 targeting to mitochondria

Several mouse models have been generated for the studies on Alsin. However, these mouse lines failed to recapitulate the phenotypes observed in human patients (Cai, Shim et al., 2008). It has recently been reported that absence of Alsin appear to specifically affect the health of corticospinal motor neurons (Gautam et al., 2016). Therefore, in order to directly probe for the role of Alsin in a more physiological background without compromising our ability for genetic and chemical manipulations, we decided to generate Alsin CRISPR knockout cells in human induced pluripotent stem cells (iPSCs). We confirmed the deletion of Alsin by both PCR and Western blot (Figure 8–figure supplement 1A,B). Importantly, we were able to further differentiate both WT and mutant (Alsin^-/-^) iPSCs to spinal motor neurons (iPSC-MNs) using a reported protocol (Reinhardt, Glatza et al., 2013). In short, we induced neural progenitor cells (NPC) through embryonic bodies formation by growing iPSC in medium supplemented with transforming growth factor-β (TGF-β) and bone morphogen protein (BMP) small molecule inhibitors (SB431542 and dorsomorphin, respectively), and WNT and Sonic Hedgehog signaling activators (CHIR99021 and PMA, respectively). Differentiation and maturation stages were achieved by culturing cells in retinoic acid (RA), cAMP, and neurotrophic factors (BDNF and GDNF) (Figure 8A). As quality control, high expression of pluripotency markers such as Oct4 and Lin28 were observed in our iPSCs as well as Nestin, Sox2 and Pax6 expression in our neuro-progenitor cells (NPCs) (Figure S8C). Differentiation into mature spinal motor neurons was validated by the expression of choline acetyltransferase (ChAT), HB9, and Islet-1 (ISL1) (Figure 8 – figure supplement 1D,E). The cells also showed extensive axonal network as revealed by MAP2 staining. Finally, mature spinal motor neurons were re-tested for expression of Alsin in both WT and Alsin^-/-^ cells (Figure 8–figure supplement 1F).

**Figure 8.**
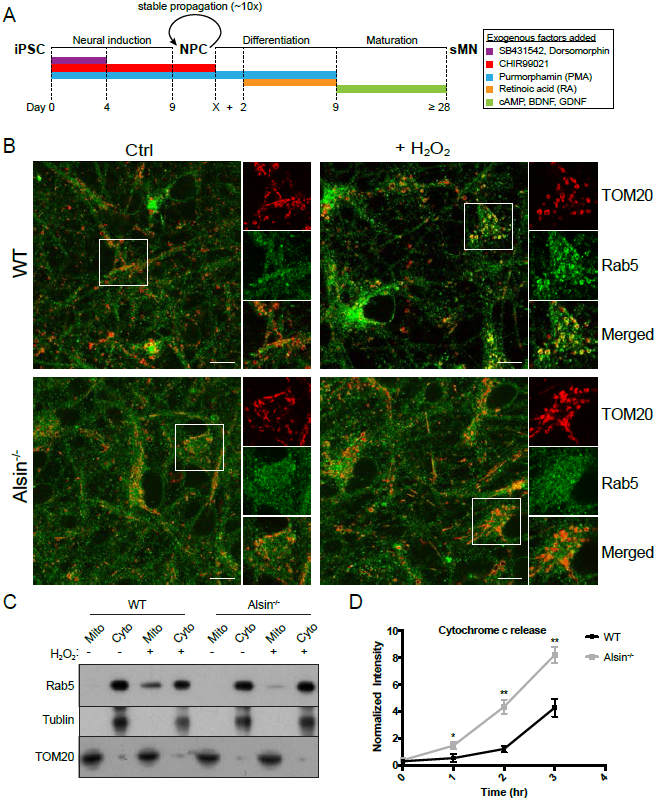
Alsin is required for Rab5 recruitment and regulates cytochrome c release. (**A**) Flow chart depicting the different stages and time (in days) from iPSC, to neuroprogenitor cells (NPC), and to generating mature spinal motor neurons (sMN). Figure legends indicate the various small molecules and compounds that are used at different stages. (**B**) WT and Alsin^-/-^ cells were challenged with 100 μM H_2_O_2_ for 1 hour. Cells were fixed and immunostained with Rab5 and TOM20 antibodies. Inset images show the representative of the signals from TOM20 and Rab5 under control (left panel) and H_2_O_2_-treated condition (right panel). Scale bars, 10 μm. (**C**) Subcellular fractionation of cytosolic (Cyto) and mitochondrial (Mito) fractions from WT and Alsin^-/-^ iPSC-sMN treated with either PBS (control) or 200 μM H_2_O_2_ for 1 hours at 37°C. Protein samples were loaded onto SDS-PAGE and imunoblotted with Rab5, tubulin (cytoplasmic loading control), and TOM20 antibodies by Western blot. (**D**) Cytosolic fractions were prepared as in Figure 6C from WT and Alsin^-/-^ iPSC-sMN challenged with 200 μM H_2_O_2_ for 1 hours at 37μC. Densitometric quantification of cytosolic cytochrome c from Western blot probed cytochrome c antibody. Data were collected from three independent experiments. Y-axis corresponds to normalized ratio intensity of experimental to loading control (* p <0.05; ** p <0.005).

We first examined the steady state localization of Rab5 and morphology of mitochondria by immunostaining for endogenous Rab5 and TOM20. In iPSC-MNs, Rab5 were present on endosomal-like vesicles ubiquitously in the soma and axon, as expected with its role in retrograde and anterograde transport (Deinhardt, Salinas et al., 2006, Guo, Farias et al., 2016). The mitochondrial network in iPSC-MNs was less tubular and consist of more numerous and smaller rounded mitochondria than those in HeLa cells (Figure 8B, Ctrl). We next wanted to verify whether iPSC-MNs would exhibit the same mitochondrial response to oxidative stress as observed in HeLa cells. Interestingly, we found that iPSC-MNs to be more susceptible to detachment and apoptosis than HeLa cells when challenged with 250 μM H_2_O_2_ for 2 h under the same conditions (data not shown). As a result, we optimized the H_2_O_2_ concentration to 100 μM for 1 hour such that no immediate cell detachment was observed during the treatment. Using a lower concentration and shorter incubation time, we then examined the morphology of mitochondria, translocation of endogenous Rab5, association of Rab5 with endosomes and mitochondria, and release of cytochrome c into cytosol. Interestingly, we did not observe significant alterations to mitochondria morphology in both WT and Alsin^-/-^ iPSC-MNs. On the other hand, WT iPSC-MNs challenged with H_2_O_2_ showed robust enrichment of Rab5 on OMM, as observed in HeLa cell, but not in Alsin^-/-^ iPSC-MNs (Figure 8B, right panel).

To further corroborate these results, we also performed subcellular fractionation in iPSC-MNs. In untreated control cells, endogenous Rab5 was present solely in the cytosolic fraction and undetectable in the purified mitochondrial fraction (Figure 8C). However, upon challenge with H_2_O_2_, Rab5 was found to co-fractionate with the mitochondrial fraction in WT iPSC-MNs but only weakly in Alsin^-/-^ iPSC-MNs (Figure 8C). The loss of Rab5 in mutant cells correlated with greater susceptibility to H_2_O_2_–induced apoptosis, as assessed by the rapid release of cytochrome c to the cytosol within an hour and further accumulation at later time points, in comparison to WT cells (Figure 8D). Collectively, our findings demonstrate that Alsin is a key regulator for recruiting Rab5 to mitochondria, which together, confer cytoprotective function against oxidative stress.

## Discussion

We discovered a novel cytoprotective mechanism during oxidative stress that entails the translocation of Rab5 from early endosomes to mitochondria. Interestingly, the activation of Rab5 requires ALS2/Alsin, that has also been implicated in early onset ALS. Our results provide an unexpected mechanistic link between the endosomal system and mitochondria that could be of primary importance for the understanding of the mechanism that cause ALS and other neurodegenerative diseases.

Different nutrient or environmental perturbations can affect mitochondria morphology and their metabolic activities such as oxidative phosphorylation and programmed cell death (Galloway & Yoon, 2013). Mitochondria can elicit responses ranging from hypoxia adaptation, inflammation, or programmed cell death when exposed to varying degrees of oxidative stress (Sena & Chandel, 2012). Our findings suggest that the endocytic system is a primary responder to mitochondria under stress. We found that laser- or exogenous ROS (e.g. H_2_O_2_)-induced damage causes MOMP, mitochondrial swelling, and release of cytochrome c, leading to caspase activation and apoptosis. In these conditions, mitochondria appear to elicit a “cue” to the endosomal system, which results in the recruitment of Alsin and subsequently Rab5 onto OMM to inhibit cytochrome c release and thus, promote cell survival (Figure 9). In the course of this study, Hammerling et al. (Hammerling, Najor et al., 2017) have reported a mitochondrial clearance mechanism by which Rab5-positive early endosomes sequester mitochondria via the ESCRT machinery when cells are treated with FCCP (carbonyl cyanide-p-trifluoromethoxypenylhydrazone), a derivative compound of CCCP. Our mechanism appears to be distinct from this and previously reported autophagic mechanisms. First, we did not observe engulfment of mitochondria into Rab5-positive early endosomes but recruitment of Alsin, Rab5 and Rabenosyn-5 on mitochondria as well as early endosome-mitochondria interactions in response to stress. Second, we did not observe such Rab5 recruitment on mitochondria in CCCP-treated cells. One explanation could be attributed to different cell types and lower concentration of CCCP employed in our experiments. Third, the recruitment of Rab5 to damaged mitochondria occurs very rapidly, i.e. within min, well preceding any autophagic components that we analyzed in this study. We found that autophagy is restricted to only a subset of small mitochondrial fragments that are LC3^+^ whereas the majority are devoid of established autophagic markers such as Parkin, LC3 and Lamp1. We cannot rule out that mitochondrial clearance mechanism may still be activated at a later time (see below). We attempted to track the fate of damaged mitochondria in a localized region after laser treatment, but the continuous photoirradation required to achieve high spatio-temporal resolution also led to quick decrease in MitoTracker Red signal and unwanted additional stress to the cell over time, which prevented us from determining its precise outcome. The loss of MitoTracker signal is likely a result of MOMP and not due to mitochondrial clearance since the OMM can be visualized by TOM20 staining and the presence of the Rab5 ring.

**Figure 9.**
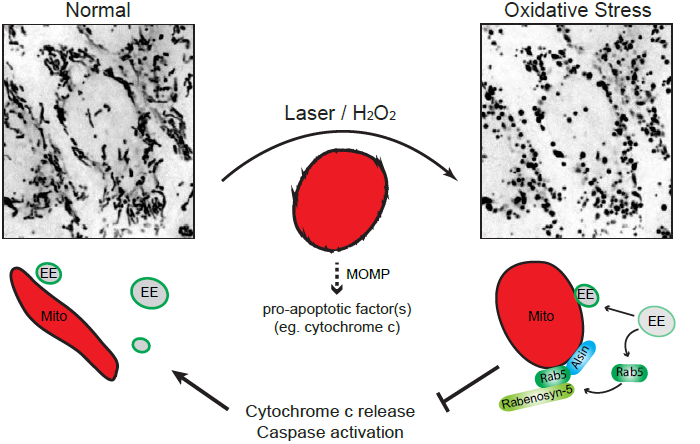
Schematic model depicting the role of Rab5-Alsin-mitochondria during oxidative stress. In normal condition, mitochondria (Mito, red) are elongated and tubular (top left). Rab5 (green) are localized on early endosomes (EE) to assemble Rab5 machinery for endosomal maturation and membrane trafficking. At steady state, we observed some EE transiently making contacts with mitochondria. During oxidative stress (eg. laser- or chemically-induced) that leads to MOMP, mitochondria undergo a dramatic morphological transformation into rounded and swollen structures (top right). This is accompanied by an increase in Rab5-positive endosomes forming MCS with mitochondria of less than <5 nm. Release of apoptotic signal such as cytochrome c from mitochondria into the cytosol is associated with Rab5 translocation to the OMM, which is involved in blocking cytochrome c release and caspase activation. The recruitment and activation of Rab5 on mitochondria depend on the Rab5 GEF Alsin (blue), resulting in a selective recruitment of Rabenosyn-5 (light green), in order to regulate apoptosis and confer cytoprotection.

Our data reveal a novel role of Alsin in the activation of the Rab5 machinery on mitochondria to regulate cell defense and survival. Alsin is a protein containing three putative GEF domains: an N-terminal regulator of chromosome condensation 1 (RCC1)-like domain (termed RLD), a middle Dbl homology/pleckstrin homology (DH/PH) domain and a C-terminal VPS9 domain. Besides studies showing the catalytic activity of Alsin VPS9 domain towards Rab5 and its role in endosomal localization and dynamics (Otomo et al., 2003, Topp et al., 2004), the physiological relevance between Alsin and Rab5 has remained mysterious. At steady state, we observe staining of Alsin on vesicular-like structures, consistent with previous reports (Millecamps, Gentil et al., 2005). In stress-induced conditions, however, Alsin and Rab5 re-localized to mitochondria. This suggests a unique mechanism leading to changes in membrane association from endosomal to mitochondrial. Our undifferentiated Alsin^-/-^ iPS cells showed a partial loss of Rab5 recruitment (data not shown) compared to mature sMN, suggesting that other GEFs may partially compensate for the loss of Alsin, depending on certain conditions. The presence of (low levels of) Rabex-5 on mitochondria (Figure 7G) suggests that this GEF might contribute to Rab5 activation, but cannot fully compensate for Alsin function. Additionally, a homologous gene encoding the protein named ALS2CL, containing just the carboxyl-terminal half of ALS2, is shown to specifically bind to Rab5 and forms a homodimer with full-length Alsin to membranous compartments (Hadano, Otomo et al., 2004, Suzuki-Utsunomiya, Hadano et al., 2007).

Which molecular mechanism is responsible for the dissociation of Rab5 from early endosomes and its recruitment to mitochondria? Several lines of evidence argue that this is a multi-step process that requires a complex cascade of factors and molecular interactions. On early endosomes, a positive feedback loop is sustained by the Rabex-5/Rabaptin-5 complex to maintain or amplify the levels of Rab5 on the membrane (Del Conte-Zerial, Brusch et al., 2008, Lippe, Miaczynska et al., 2001, Zhang, Zhang et al., 2014). However, the stress response introduces instability to such a system. Activation of p38 MAPK by H_2_O_2_ has been shown to stimulate formation of the GDI:Rab5 complex, thus extracting Rab5 from the early endosome membrane (Cavalli, Vilbois et al., 2001). It is conceivable that such mechanism may account for the mobilization of Rab5 from the endosomal membrane. In addition, p38 MAPK modulates endosomal function via phosphorylation and membrane association of Rab5 effectors (Mace, Miaczynska et al., 2005). We found a reduction in the levels of Rab5 co-localizing with its endosomal effectors (eg. EEA1) (Figure 7D). This may provide an energy-sparing mechanism that overrides Rab5 endosomal interactions in order to slow down endocytosis by relocating part of the Rab5 machinery to mitochondria. Consistent with this idea, Rabaptin-5 is shown to be selectively cleaved by capase-3 during apoptosis, thus affecting its interactions with Rab5 and reducing overall endocytic capacity (Cosulich, Horiuchi et al., 1997, Swanton, Bishop et al., 1999). The loss of endocytic capacity is supported by the fact that we observed a severe block in transferrin uptake. Such a block may confer another cytoprotective role by reducing iron uptake in order to avoid iron overload and toxicity, which is often observed in neurodegeneration (Nunez, Urrutia et al., 2012). Interestingly, hippocampal HT-22 neurons exposed to excess iron trigger mitochondrial fragmentation and result in decreased cell viability (Park, Lee et al., 2015). As for the activation of Rab5 on mitochondria, the N-terminal RLD of Alsin has been shown to exhibit an autoinhibitory effect on its VPS9 domain (Otomo et al., 2003). We posit that mitochondrial-induced stress triggers structural changes in the protein, releasing the autoinhibitory effect of RLD, thereby exposing the VSP9 domain for Rab5 activation and recruitment.

Besides the decrease in iron uptake to reduce free radical toxicity, a key question is what is the functional significance of the assembly of the Rab5 machinery on mitochondria? It appears that the stress response triggers the remodeling of the OMM to confer molecular features characteristic of the endocytic system. The Rab5 machinery may be used to bring mitochondria in close proximity to early endosomes to form MCS, as these appear to increase upon stress. These MCS may mediate the transfer of lipids and metabolites required for the repair response (Helle, Kanfer et al., 2013). However, the Rab5 translocation may be a priming step of a stress response pathway that could subject the mitochondria to interactions with the entire endo-lysosomal system, eventually leading to autophagy or apoptosis. One quality control mechanism is the formation of mitochondrial-derived vesicles (MDVs), which are involved in the transport of oxidized or damaged cargo to late endosomes and lysosomes for degradation (Soubannier, McLelland et al., 2012). This process primarily depends on PINK1/Parkin (McLelland, Soubannier et al., 2014) but can also occur in a PINK1/Parkin-independent manner (Matheoud, Sugiura et al., 2016). Rab5 could play a role in MDV formation although we could not detect vesicle budding events within our time frame and experimental conditions. Once recruited onto mitochondria, Rab5 activity may not be limited to the recruitment of its effectors but initiate a more extensive endosomal Rab cascade via the Rab coupling/conversion mechanism. On early endosomes, Rab5 interacts with divalent effectors coupling its activity to Rab proteins (e.g. Rab4, Rab11) required for receptor recycling (de Renzis, Sonnichsen et al., 2002, Vitale, Rybin et al., 1998). Rab5 also initiates activation of Rab7 resulting in conversion of early endosomes into late endosomes (Rink et al., 2005), and consistent with this, silencing of Rab5 in mouse liver causes the loss of the entire endolysosomal system (Zeigerer et al., 2012). Rab coupling/conversion may thus be initiated also on the mitochondria and therefore, it is possible that the mitochondria-endosome MCS may evolve with time leading to the engulfment of mitochondria by the early endosomes (Hammerling, Najor et al., 2017) or to the conventional autophagic processes (Ao, Zou et al., 2014, Stolz, Ernst et al., 2014). It will therefore be important to explore the dynamics of other endosomal Rab GTPases in relation to Rab5 over time.

The physiological role of Alsin has been linked to both endosomes and mitochondria. Cultured hippocampal neurons from Alsin knockout mice display an accumulation of enlarged Rab5 endosomes and reduced endosomal motility (Lai, Xie et al., 2009). Mutational and linkage analysis of Alsin from human patients show that VPS9 domain is critical for Alsin function (Daud, Kakar et al., 2016, Verschuuren-Bemelmans, Winter et al., 2008). Recent EM study on corticospinal motor neurons (CSMN) from Alsin KO mice reveals a selective defect in mitochondria morphology displaying defective membrane and vacuolated cristae (Gautam et al., 2016). Interestingly, WT vs Alsin KO CSMN shows no change in Parkin expression, suggesting that mitophagy does not play a major role. We postulate that the pathological condition of mitochondrial defects in Alsin KO cells is related to a deficiency in Rab5 recruitment to mitochondria, thereby leading to low-levels of protection from ROS accompanying aging. In ALS patients, motor neurons accumulate likely more damaged mitochondria as they age, which eventually become an overload for cells.

The cause for ALS is still not fully understood, but oxidative stress is considered to be a major contributor. Mutations in the anti-oxidant enzyme, superoxide dismutatse 1 (SOD1), are associated with motor neuron degeneration. Mouse model shows that an accumulation of the SOD1 mutant proteins results in mitochondrial swelling and increased oxidative damage (Jaarsma, Rognoni et al., 2001). Interestingly, loss of Alsin in the mutant SOD1 transgenic mice exacerbates and accelerates disease progression (Hadano et al., 2010). These studies, along with our findings, corroborate the protective role of Alsin during oxidative stress. The mechanistic link between Rab5 and Alsin may present a general or related mechanism in other neurodegenerative diseases. In Parkinson disease, the most common mutation found in the multidomain Leucine-rich repeat kinase 2 (LRRK2) protein leads to hyper-activation of the kinase domain, resulting in hyper-phosphorylation of a number of Rab GTPase substrates including Rab5 (Steger, Tonelli et al., 2016). This may present yet another mechanism of regulating Rab5 localization and function on mitochondria. Future work using different neurodegenerative disease models in differentiated human neurons will provide deeper insights into the disease etiology.

## Materials and Methods

### Cell lines, cell culture, and growth conditions

HeLa cells were cultured in high glucose DMEM (Gibco) with 10% fetal bovine serum, 100 U/ml penicillin, 100 μg/ml streptomycin, and 2 mM glutamine (all reagents from Sigma-Aldrich) at 37°C with 5% CO_2_. All plasmids were transfected using Effectene transfection reagent (Qiagen) according to manufacturer’s protocol. All bacterial artificial chromosome (BAC) transgene HeLa cell lines expressing different markers were obtained from the BAC recombineering facility at MPI-CBG (Dresden, Germany) and generated using the methods previously described (Poser, Sarov et al., 2008).

### Plasmids and chemical reagents

Construction of the pEGFP-C3-2xFYVE was constructed using mouse Hrs FYVE domain containing a linker (QGQGS) (Raiborg, Bremnes et al., 2001). Alexa-conjugated transferrin (Invitrogen; T13342) and EGF (Invitrogen; E13345) were used at 25 μg/ml and 2 μg/ml, respectively. Carbonyl cyanide 3-chlorophenylhydrazone (CCCP) was purchased from Sigma Aldrich (C2759). Stock solution was made to the final concentration of 10 mM in DMSO. Hydrogen peroxide (H_2_O_2_) (Merck Millipore; 7722-84-1) 100 mM stock solution was prepared in PBS.

### Live-cell confocal imaging

Cells were plated in a 35-mm petri dish, 14-mm glass bottom microwell for live-cell imaging. Before imaging, medium was replaced with HEPES-buffered DMEM without phenol red (Gibco). Cells were then imaged by time-lapse microscopy (Spinning Disc, Andor-Olympus-IX71 inverted stand microscope and Nikon TiE inverted stand microscope equipped with spinning disc scan head (CSU-X1; Yokogawa), fast piezo objective z-positioner [Physik Instrumente], and back-illuminated EMCCD camera (iXon EM+ DU-897 BV; Andor). Imaging was done with an Olympus UPlanSApo 100x 1.4 Oil and Nikon Apo 100x 1.49 Oil DIC 0.13–0.20 objectives (illumination by lasers: DPSS-488nm, DPSS-561nm, DPSS-640nm). Individual planes were recorded at ∼10 frames per sec with Z-stacks of three planes (step 0.3 μm).

### Photosensitization of mitochondria

Cells were incubated with MitoTracker Red CMXRos (ThermoFisher; M7512) at a final concentration of 100 nM for 30 min at 37°C, 5% CO_2_ incubator, and followed by 2X PBS wash before irradiating with 561 nm laser on the spinning disc Andor-Olympus-IX71 at low power dosage of ∼5 J/cm^2^ for 60 seconds.

### Correlative light electron microscopy

Cells were grown on gridded dish (ibidi μ-Dish 35-mm, high Grid-500). Cells in different locations were laser-treated with 561 nm laser for 30 secs. Cells were fixed in 2.5% glutaraldehyde/PBS for 30 min at room temperature. Post-fixation and embedding were performed using 1% osmium tetroxide/1.5% potassium ferrocyanide and Epon Lx112, respectively. Sectioning of 150 nm thick UA sections was performed on a Leica Ultracut UCT (Leica Microsystem, Wetzlar, Germany) with a diamond knife. Samples are post-stained with 2% uranyl acetate and lead citrate. 2D images were acquired on a Tecnai T12 (FEI, Hillsboro, Oregon, USA).

### Immunofluorescence and antibodies

Cells were seeded on ibidi Grid-500 glass bottom. After laser or H_2_O_2_ treatment, cells were fixed in 4% paraformaldehyde/PBS for 15 min at room temperature. Cells were washed twice with PBS and permeabilized in PBS containing 0.1% saponin, and 1% BSA for 30 min at room temperature. Cells were immunostained with corresponding primary: anti-rabbit Rabenosyn-5/ZFYVE20 (Sigma Aldrich: HPA044878), anti-mouse EEA1 (BD Biosciences: 610457), anti-rabbit TOM20 (Santa Cruz Biotechnology: sc-11415), anti-rabbit APPL1 (Abcam: ab59592), anti-mouse Rab5 (BD Biosciences: 610724), anti-mouse cytochrome c (Abcam: ab6311), and anti-rabbit Alsin (Novus Biological: NBP2-14284) antibodies. Alexa fluor-conjugated from ThermoFisher were used as secondary antibodies. Samples were mounted with Mowiol (Sigma-Aldrich) on glass slides and examined using the Zeiss LSM 880 inverted single photon point scanning confocal system with Quasar detector (32 spectral detection channels in the GaAsP detector plus 2PMTs) and transmitted light detector. Acquired images were processed and saved using the Zeiss ZEN software. For immunofluorescence on iPSCs, smNPCs, and sMNs, cells were fixed with 4 % formaldehyde for 10 min, washed three times with wash buffer (0.3 % Triton-X in PBS) for 5-10 min, and blocked with blocking buffer (5 % goat serum, 2 % BSA, and 0.3 % Triton-X in PBS) for 1 hour at room temperature. Cells were incubated with primary antibody in blocking buffer overnight at 4 °C. After washing three times with PBS for 10 min, cells were incubated with secondary antibodies in wash buffer for 2-3 hours at room temperature followed by three washes in PBS for 10 min. Primary antibodies: goat anti-ChAT (1:100) (Millipore, #AB144P), mouse anti-HB9 (1:50) (DSHB, #81.5C10, conc.), rabbit anti-ISL1 (1:100) (Abcam, #ab20670), mouse anti-LIN28 (1:1000) (Cell signaling, #5930S), chicken anti-MAP2 (1:1000) (Novus Biologicals, #NB300-213), mouse anti-Nestin (1:150) (R&D Systems, #MAB1259), rabbit anti-OCT4 (1:500) (Abcam, #ab19857), rabbit anti-PAX6 (1:300) (Covance, #PRB-278P), and rabbit anti-SOX2 (1:500) (Abcam, #ab97959).

### Transferrin uptake

Cells were seeded in a 384-well plate and incubated with either complete medium or in the presence of 250 μM H_2_O_2_ for 2 hours at 37°C. Cells were then pulsed with Alexa-647 Tfn (10 μg/ml) for 5 min or 10 min, followed by 3x PBS wash, fixed with 3.7% PFA for 15 min, and then stained with DAPI (1:1000) and CellMask Blue (1:2000) (ThermoFisher). Image acquisition was performed via the automated confocal imaging system, CV7000S Yogokawa. Images analysis were performed using MotionTracking software.

### Subcellular fractionation

Cytosolic and mitochondrial fractions were performed using the mitochondria isolation kit, according to manufacturer’s protocol with minor modification (ThermoFisher: cat89874). Cells (∼1 x 10^7^) were resuspended in 400 μl Mitochondrial Isolation Reagent A. Cells were chemically lysed by adding 5 μl of Reagent B. After 5 min incubation on ice, 400 μl of Reagent C was added to each sample and centrifuged at 720 x g for 10 min. Supernatant was transferred to a new eppendorf tube and centrifuged at 3000 x g for 15 min at 4°C. Supernatant was collected and trichloroacetic acid (TCA)/acetone precipitation was performed to obtain the final cytosolic fraction. The remaining pellet was washed by adding 500 μl of Reagent C and centrifuged at 15,000 x g for 5 min. Final samples were resuspended in SDS loading buffer.

### Cytochrome c release assay and Western blot

Cells were seeded onto a 12-well plate. For hydrogen peroxide treatment, reagent was added directly into the well to achieve the appropriate concentration. Separation of mitochondrial and cytosolic fractions was performed using mitochondrial isolation kit from ThermoFisher (cat:89874) with an additional step of trichloroactic acid precipitation of the cytosolic fraction. The final pellet was dried for 2-3 min in a 95 °C heat block before resuspending it in SDS loading buffer. Cell lysates were separated by SDS-PAGE, transferred onto nitrocellulose membrane and blocked in 5% milk in PBS containing 0.1% Tween. Primary and secondary antibodies were diluted in blocking buffer and incubated for 2 h at room temperature. Detection of bands was performed using electrochemiluminescence reagent and exposure onto x-ray films. The following antibodies were used by Western blot: anti-mouse cytochrome c (Abcam: ab13575), anti-rabbit gamma tubulin (Sigma-Aldrich: T6557), anti-rabbit Alsin (Sigma Aldrich: SAB4200137), anti-mouse EEA1 (BD Biosciences: 610457), and anti-rabbit TOM20 (Santa Cruz Biotechnology: sc-11415).

### Generation of CRISPR/Cas9 knockout in human induced pluripotent cells

Human KOLF_C1 iPSC (kindly provided by Bill Skarnes, Sanger Institute) were cultured in feeder-free conditions on Matrigel with TeSR-E8 media (StemCell, Germany). For ALS2/Alsin knockout using CRISPR/Cas9 genome editing, 350,000 cells were detached using Accutase, washed once with PBS and electroporated using the Neon Transfection System (Invitrogen, Germany, 10ul kit, 1000V, 20ms, 3 pulses). Genomic sequence of human Alsin was analysed for CRISPR/Cas9 target sites by Geneious 8.1.6 software (Biomatters), and two pairs of guides flanking a critical exon (exon3) were selected (5’-GCTAAAGTACTGAATTTTGG-3’ and 5’-AATAAAATCAGCAGGTGTGG-3’; 5’-GAATTTCTACAAAGTGCAGG-3’ and 5’-TAGCCTGGATGATGGCCGTT-3’) and were used together to cause a frame shifting exon deletion). The in vitro efficiency of these gRNAs was assessed by generating genomic PCR cleavage template of 3.4 bp (primers used: for-CCTCCCTTCCCAGGATCTGA and rev-TGCTCAACTCGAGTGCCTTT; for-CAGGGTGAGCATCCCACATT and rev-AGGAGTTCCAGTCAACCAGT) and incubating with recombinant Cas9. All gRNAs used in vitro were identical in sequence to the DNA sense strand and not complementary to the mRNA sequence. The RNAs employed in this method were chemically-modified and length optimized variants of the native guide RNAs (Alt-R^TM^ CRISPR crRNAs and tracrRNA, Integrated DNA Technologies, Coralville, IA, USA). Recombinant Cas9 (provided by Protein Expression Facility at MPI-CBG) protein from *Streptococcus pyogenes* was used. The crRNAs were mixed with trRNA and NLS-Cas9 (1 μg/μl). The guide RNA complex was formed by mixing the crRNAs and tracrRNAs in equal amounts in Buffer R (Invitrogen, Germany) at 100μM concentration. 5 days after electroporation, cells were pooled and seeded for clonal dilution. Single clones were mechanically picked and amplified. Next, genomic DNA had been isolated using QuickExtract DNA Extraction Solution (EpiCentre, USA). Homozygous deletions had been verified by PCR and sequencing.

### Generation of iPSC-derived smNPC, and differentiation of smNPCs to MNs

All procedures were performed as previously described (Reinhardt et al., 2013). Briefly, for smNPC generation, iPSC colonies detached from Matrigel-coated wells by 1 mg/ml dispase were resuspended in hESC medium (DMEM/F12, 20 % KnockOUT Serum Replacement, 1 % Penicillin/Streptomycin/Glutamine, 0.1 mM Non-Essential Amino Acids Solution, 0.05 mM beta-mercaptoethanol, without bFGF) supplemented with 10 μM SB431542 (Tocris, #1614), 1 μM dorsomorphin (Tocris, #3093), 3 μM CHIR99021 (Axon Medchem, #Axon-1386) and 0.5 μM purmorphamine (STEMCELL Technologies, #72202), and cultured in non-coated petri dishes. After two days, hESC medium was replaced by N2B27 medium (1:1 mixture of DMEM/F12 and Neurobasal medium, 1 % Penicillin/Streptomycin/Glutamine, 1:100 B-27 supplement minus vitamin A, 1:200 N-2 supplement) supplemented with the same small molecules as listed above. After another two days, culture medium was replaced by smNPC expansion medium (N2B27 medium supplemented with 150 μM ascorbic acid (Sigma, #A4403), 3 μM CHIR99021 and 0.5 μM purmorphamine). At day 6 of neural induction, embryonic bodies were broken into smaller clumps by titration and plated in 6 wells of a Matrigel-coated 12-well plate. At day 9, cells were passaged for the first time using Accutase at a 1:3 split ratio and seeded in 4 wells of a Matrigel-coated 6-well plate. Afterwards, cells were passaged ones a week and seeded at a density of 1 x 10^6^ cells per well. To obtain a highly pure smNPC culture, smNPCs were propagated for at least 10 passages in smNPC expansion medium. For differentiation of smNPC to MNs, smNPCs were seeded at a density of 1.5 x 10^6^ cells per one well of a Matrigel-coated 6-well plate and cultured in N2B27 medium supplemented with 1 μM purmorphamine for the first two days of differentiation. The cells were then cultured in N2B27 medium supplemented with 1 μM purmorphamine and 1 μM retinoic acid (Sigma, #R2625) until day 9 of differentiation. At day 9, cells were dissociated using Accutase and plated on polyornithine/laminin-coated ibidi μ-slides (at a density of 150000 cells per well) or Nunc 4-well plates (at a density of 300.000 cells per well) in maturation medium (N2B27 medium supplemented with 0.5 mM cAMP (Sigma, #D0627), 10 ng/ml BDNF (Peprotech, #450-02-10), and 10 ng/ml GDNF (Peprotech, #450-10-10)). Cells were maintained in maturation medium until analysis on day 28.

### Image and statistical analysis

Image resizing, cropping and brightness were uniformly adjusted in Fiji (<u>http://fiji.sc/</u>). Co-localization analysis was performed using MotionTracking software (Rink et al., 2005) (<u>http://motiontracking.mpi-cbg.de/get/</u>) and described previously (Gilleron, Querbes et al., 2013). The y-axis is expressed as the ratio of co-localized objects (eg. A to B) to total objects found in A. Final images were assembled using Adobe Photoshop and Illustrator. Densitometry quantification were performed in Fiji following the previously described protocol (<u>http://www.yorku.ca/yisheng/Internal/Protocols/ImageJ.pdf)</u>.

## Acknowledgements

We thank the MPI-CBG light microscopy facility for access and technical assistance; the TransgeneOmics facility and Hyman lab, especially Mihail Sarov, Aleksandra Syta, Christina Eugster, Kathleen Rönsch, Marit Leuschner, and Ina Poser, for the design, generation, and maintenance of the CRISPR/Cas9 KO iPSC lines; Julia Japtok from the lab of Andreas Hermann for sharing protocols for the smNPC generation and MN differentiation; Weihua Leng for technical assistance and method discussion for electron microscopy; Ina Nuesslein and Christina Eugster from the MPI-CBG FACS facility for assistance on the flow cytometry; Rico Barsacchi from the Technology Development Studio facility for assistance with the transferrin uptake and access to the Yokogawa system; Yannis Kalaidzidis for assistance with the MotionTracking software; and Heidi McBride for discussion and feedbacks.

---

**Figure 1–figure supplement 1.**
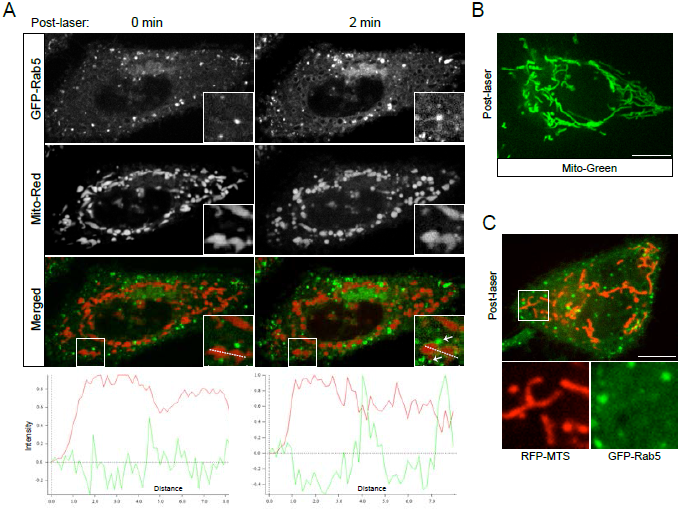
Rab5 recruitment to mitochondria occurs within min and is specific to MitoTracker-Red CMXRos. (**A**) HeLa BAC GFP-Rab5 cells labeled with MitoTracker-Red CMXRos were photoirradiated with low dosage of 561 nm laser (∼5 J/cm^2^) for 60 sec. Snapshot of a cell was taken immediately after photoirradiation (0 min) and 2 min elapsed. Inset images show the rapid recruitment of Rab5 signal (green) around mitochondria (red) via line trace (red: Mito-Red; green: GFP-Rab5). Arrowheads indicate endosomes that are in close proximity to mitochondria. (**B**) HeLa cells were labeled with MitoTracker-Green FM (Mito-Green) and photoirradiated with 561 nm laser (∼10 J/cm^2^) for 60 sec. Image was taken 20 min post-laser. Mitochondrial network remained tubular. (**C**) HeLa BAC GFP-Rab5 cells were transfected with TagRFP-MTS and photoirradiated with 561 nm laser (∼10 J/cm^2^) for 60 sec. Image was taken 20 min post-laser. Scale bars, 10 μm.

**Figure 1–figure supplement 2.**
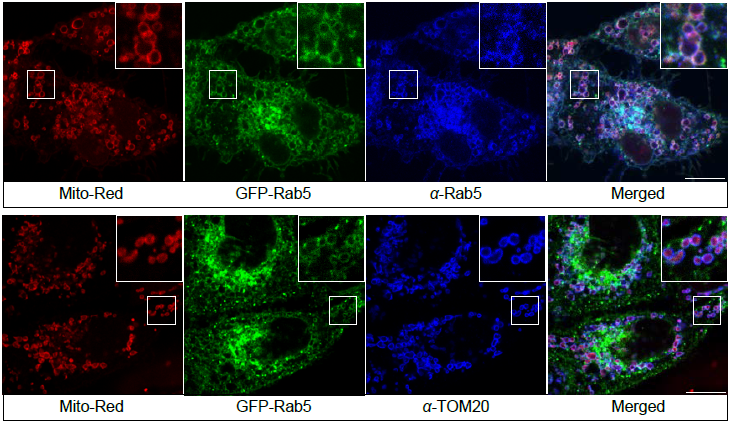
Localization of GFP-Rab5 signals assessed by Rab5 and TOM20 antibodies. BAC GFP-Rab5 cells were seeded on gridded glass bottom, labeled with MitoTracker-Red CMXRos, photoirradiated as before, and fixed after 30 min post-laser. Cells were then immunostained with specific antibodies against Rab5 or TOM20. Scale bars, 10 μm.

**Figure 4–figure supplement 1.**
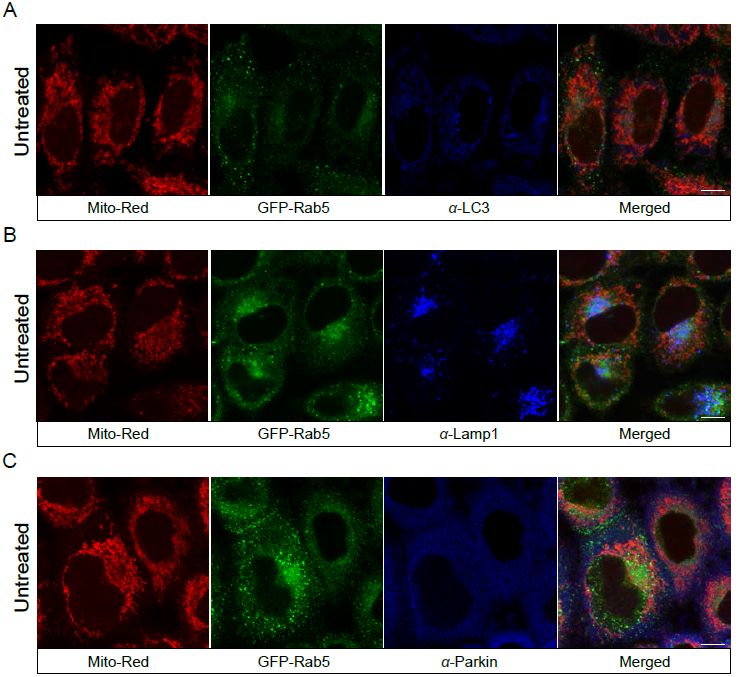
Localization of endogenous LC3, Lamp1, and Parkin in untreated cells. BAC GFP-Rab5 cells were seeded on gridded glass bottom, and labeled with MitoTracker-Red CMXRox. Fixed cells were immunostained with specific antibodies against LC3, Lamp1, and Parkin. Cells shown were imaged from area outside of laser-treated cells. Scale bars, 10 μm.

**Figure 4–figure supplement 2.**
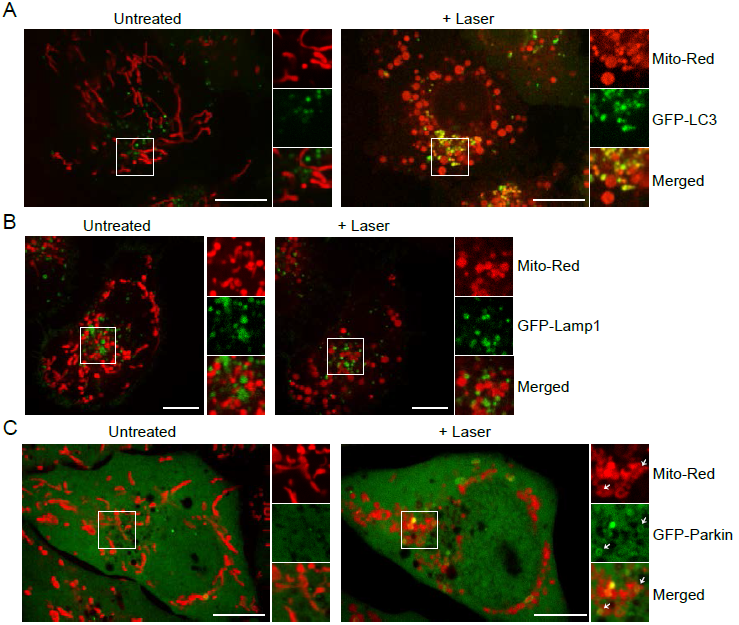
BAC GFP-LC3, GFP-Lamp1, and GFP-Parkin localization upon laser-induced stress. Live-cell imaging of different BAC cell lines. Cells were labeled with MitoTracker-Red CMXRos and photoirradiated as before. Images of the same cell in pre-(untreated) and post-laser are shown. Inset images indicate the approximate area before and after laser treatment. (**A**) Images of BAC GFP-LC3 cell before and 60 min post-laser. (**B**) Images of BAC GFP-Lamp1 cell before and 60 min post-laser. (**C**) Images of BAC GFP-Parkin cell before and 60 min post-laser. Scale bars, 10 μm.

**Figure 7–figure supplement 1.**
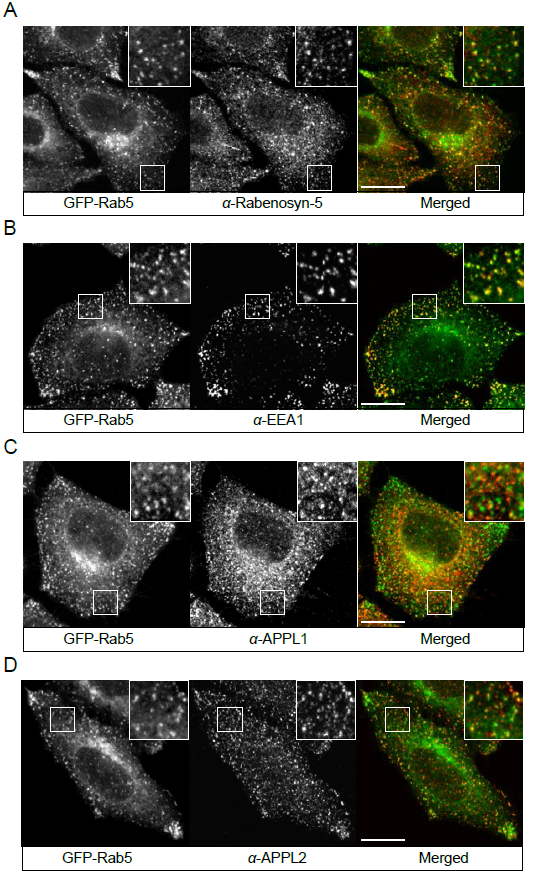
Localization of endogenous Rabenosyn-5, EEA1, and APPL1/2. BAC GFP-Rab5 cells were seeded on glass coverslip, fixed, and immunostained with specific antibodies against Rabenosyn-5 (**A**), EEA1 (**B**), APPL1 (**C**), and APPL2 (**D**). Scale bars, 10 μm.

**Figure 7–figure supplement 2.**
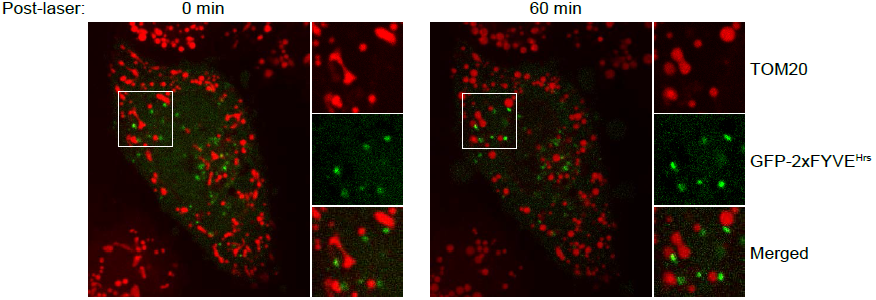
HeLa cells over-expressing GFP-2xFYVE^Hrs^. Cells were transfected with GFP-2xFYVE^Hrs^ for 24 h, labeled with MitoTracker-Red CMXRos, photoirradiated as before, and then imaged live. Images were capture immediately post-laser (0 min) and after 60 min elapsed time.

**Figure 7–figure supplement 3.**
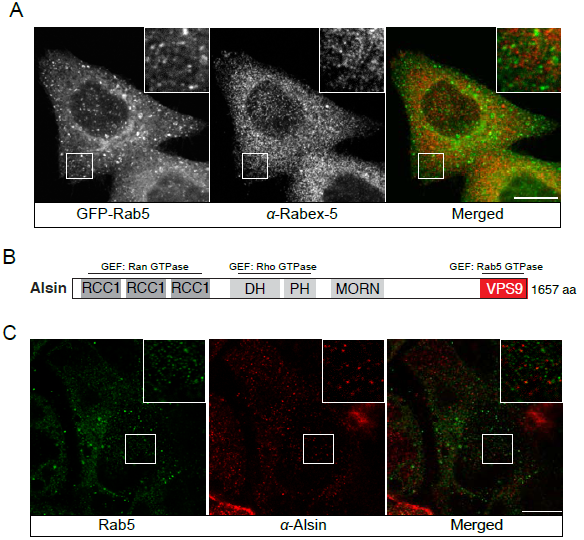
Localization of Rabex-5 and Alsin in HeLa cells. (**A**) BAC GFP-Rab5 cells were seeded on glass coverslip, fixed, and immunostained with specific antibodies against Rabex-5. (**B**) A schematic representation of domain organization of full-length human Alsin. The regulator of chromosome condensation 1 (RCC1), B-cell lymphoma (Dbl) homology (DH) and pleckstrin homology (PH), membrane occupation and recognition nexus (MORN) motif, and vacuolar protein sorting 9 (VPS9) domains are labeled. Three putative GEF domains are annotated. (**C**) Immunostaining of HeLa cells with specific antibodies against Rab5 and Alsin.

**Figure 8–figure supplement 1.**
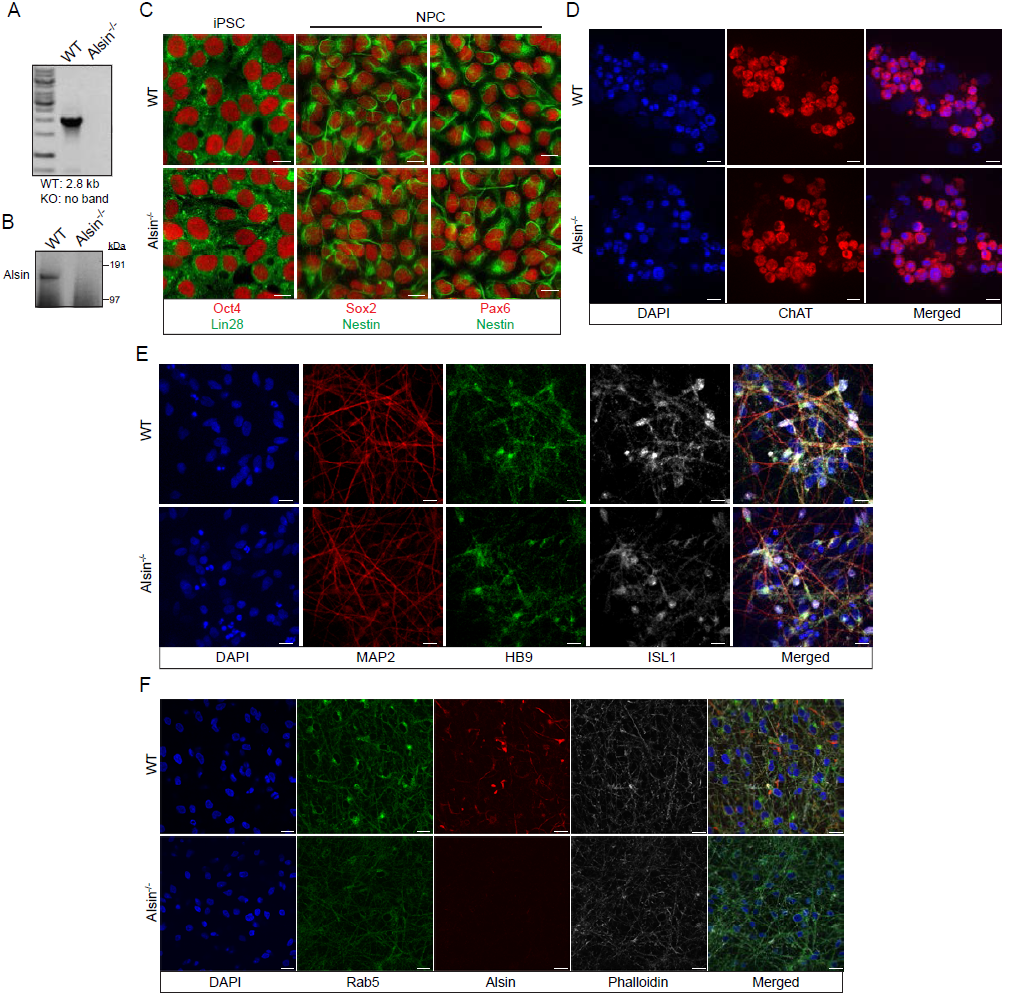
Validation of CRISPR/Cas9 Alsin^-/-^ cells in induced pluripotent stem cells (iPSC) and iPSC-derived spinal motor neurons (iPSC-sMN). (**A**) Electrophoresis result of the PCR reaction using primers flanking exon 3 and within exon 3. Homozygous deletion was confirmed by the absence of a 2.8 kb in Alsin^-/-^. (**B**) Protein lysates from WT KOLF and Alsin^-/-^ iPSC were loaded onto SDS-PAGE and probed with antibody against Alsin. A band detected at ∼184 kDa, but absent in Alsin^-/-^, corresponds to full-length Alsin. (**C**) WT KOLF and Alsin^-/-^ iPSC and neuroprogenitor cells (NPC) were immunostained with pluripotency markers Oct4 and Lin28, and neuroprogenitor markers Sox2 and Pax6, respectively. (**D-E**) WT KOLF and Alsin^-/-^ iPSC-sMN were immunostained with motor neuron markers such as ChAT, HB9, and ISL1, nuclear dye DAPI, and cytoskeletal marker MAP2. (**F**) WT KOLF and Alsin^-/-^ iPSC-sMN were immunostained with specific antibodies against Rab5 and Alsin, along with DAPI (nuclear) and phalloidin (actin) stains. Scale bars, 10 μm.

**Video 1. Dynamics of transferrin and mitochondria.** HeLa cells were labeled with Alexa-488 transferrin (Tf; green) and MitoTracker-Red (Mito; red). Images were acquired using a spinning disk confocal microscope at 11 frames/sec. Time stamp corresponds to min:sec:msec.

**Video 2. Dynamics of transferrin and mitochondria.** Zoom-in of a HeLa cell labeled with Alexa-488 transferrin (Tf; green) and MitoTracker-Red CMXRos (Mito; red). Movement of endosomes during a mitochondria fission event is shown. Images were acquired using a spinning disk confocal microscope at 11 frames/sec. Time stamp corresponds to min:sec:msec.

**Video 3. Dynamics of Rab5-positive endosomes and mitochondria.** Zoom-in of a BAC GFP-Rab5 (green) cell labeled with MitoTracker-Red CMXRos (Mito; red). Images were acquired using a spinning disk confocal microscope at 11 frames/sec. Time stamp corresponds to min:sec:msec.

**Video 4. Mitochondria dynamics during laser-induced stress.** HeLa cells were labeled with MitoTracker-Red CMXRos and photoirradiated via 561 nm laser for 1 min, and immediately imaged. Time-lapse was acquired using a spinning disk confocal microscope at 11 frames/sec for 5 min.

**Video 5. Endosomes and mitochondria interactions upon laser-induced stress.** BAC GFP-Rab5 (green) cells were labeled with MitoTracker-Red CMXRos (red) and photoirradiated via 561 nm laser for 1 min, and then imaged after 5 min. Time-lapse was acquired using a spinning disk confocal microscope for 20 frames with 5 sec increment. Boxed regions indicate distinctive GFP-Rab5 endosomes docking onto mitochondria.

**Video 6. Mitochondria dynamics during H_2_O_2_-induced stress.** HeLa cells were labeled with MitoTracker-Red CMXRos and then imaged in the presence of 250 μM H_2_O_2_ for 60 min. Time-lapse was acquired using a spinning disk confocal microscope for 50 frames with 2 min increment.

**Figure 1–Source Data.** Numerical data corresponding to the graph presented in Figure 1D.

**Figure 4–Source Data.** Numerical data corresponding to the graph presented in Figure 4G.

**Figure 5–Source Data.** Numerical data corresponding to the graphs presented in Figure 5C,D.

**Figure 6–Source Data.** Numerical data corresponding to the graphs presented in Figure 6C,E.

**Figure 7–Source Data.** Numerical data corresponding to the graphs presented in Figure 7C,D,G.

**Figure 8–Source Data.** Numerical data corresponding to the graph presented in Figure 8D.

